# TCR-dependent and TCR-independent *in-vitro* T cell activation generate distinct functional, metabolic, and cytokine programs: Protein kinase C signalling augments anti-CD3+anti-CD28 responses

**DOI:** 10.64898/2026.07.15.738657

**Authors:** Nikita S. Ramteke, Dipankar Nandi

**Author notes:** Corresponding author: FE-14, Department of Biochemistry, Biological Sciences Building, C.V. Raman Road, Indian Institute of Science, Bengaluru-560012, Karnataka, India.

## Abstract

**Introduction:** T cell activation is central to the adaptive immune response. *In vitro* studies on T cell activation often utilize two distinct approaches: first, engaging T cell receptors (TCR) using plate-bound αCD3 together with soluble αCD28 (TCR-dependent). Second, triggering intracellular signalling cascades using phorbol 12-myristate 13-acetate (PMA) and Ionomycin or P+I (TCR-independent). Both methods are widely used; however, a systematic comparison of the activation methods across a range of stimulation strengths to evaluate their effects on T cell function and metabolism has not been investigated in great detail. In this study, we compared the consequences of engaging T cells using TCR-dependent and TCR-independent activation pathways across varying signal strengths.

**Methods:** T cells from BALB/c mice were isolated and activated under four conditions: αCD3, αCD3+αCD28, PMA with low Ionomycin (P+IL) and PMA with high Ionomycin (P+IH). We studied differences with respect to several parameters: morphology, flow analysis, metabolic activities, cytokines. The roles of Protein kinase C (PKC) and Ca²⁺ pathways were addressed by supplementing αCD3+αCD28 cultures with different doses of exogenous PMA or Ionomycin.

**Results:** P+I activation outperformed the αCD3+αCD28 activation system across most readouts by displaying enhanced blasts, higher cycling, greater glucose uptake, increased lactate and ROS production, together with higher upregulation of CD25 and CD44 activation markers. P+IH activation dampened several responses including CD69 expression. CD4 co-receptor was downregulated greatly with P+I activation but not αCD3+αCD28. Most cytokines followed signal strength comparably between both systems; however, differences were observed with others: P+I stimulation favoured IL-6 and IL-12 induction whereas αCD3+αCD28 activation preferentially induced CCL2 and IL-1β. Importantly, PKC activity was substantially lower upon αCD3+αCD28 stimulation and the addition of PMA, but not Ionomycin, to αCD3+αCD28 cultures enhanced proliferation, metabolism and expression of activation markers.

**Discussion:** TCR-dependent and TCR-independent T cell activation models have clear functional and metabolic differences. The observation that PKC signalling can boost T cell activation with αCD3+αCD28 is likely to be significant and may have translational implications such as CAR-T cell anti-tumor therapy where αCD3+αCD28 stimulation is widely used. The implications of our findings with regard to augmenting T cell mediated immunotherapies are discussed.

## Introduction

The immune system has evolved a flexible and dynamic strategy for mounting specific responses against a diverse range of antigens. Central to this process are T cells, the key cellular mediators of adaptive immunity, which are activated upon exposure to antigen presented by antigen-presenting cells (APCs). When activated, naïve T cells exit the G0 phase of the cell cycle and undergo a series of signalling events that drive their proliferation, differentiation and effector functions. Activation of T cells is a consequence of multiple ligand-receptor interactions that occur at the interface of a T cell and an APC. Naive T cell activation requires two main signals. The first signal initiates with engagement of the TCR by a short peptide antigen fragment bound to a major histocompatibility complex (MHC). The antigen induced stimulation of T cell receptor (TCR) delivers the primary signal and the second signal in initiating activation and co-stimulation is via CD28 binding to CD80/CD86 present on APCs which leads to effector response that includes cytokine production and cytotoxic killing (Linsley and Ledbetter, 1993).

Engagement of the TCR, together with costimulatory receptors such as CD28, sets off intracellular signalling cascades that rely on second messengers to relay and shape the response. Accessory molecules fine-tune this process at several levels: adhesion molecules like LFA-1 and CD2 stabilise the contact between the T cell and the APC and co-receptors such as CD4 and CD8 modify the signal delivered through the TCR. These signalling events occur over different timescales. Early events, including protein phosphorylation, ion fluxes, and changes in gene transcription, occur within minutes to hours. Later responses develop over several days, collectively drive T cell proliferation, differentiation, and effector function (Smith-Garvin, Koretzky and Jordan, 2009). T cell activation leads to activation of further downstream signaling events which include increase in intracellular Ca^2+^, activation of phospholipase C-γ1 (PLCγ1), generation of the secondary messengers diacylglycerol (DAG) and inositol-1,4,5-trisphosphate (IP3), recruitment and activation of PKC-θ and these signals together activates the transcription factors NF-κB, AP-1, and NFAT leading to clonal expansion and differentiation of T cells (Isakov and Altman, 2002; Smith-Garvin, Koretzky and Jordan, 2009; Brownlie and Zamoyska, 2013).

Sub-optimal activation of T cells leads to inefficient expansion of T cells (Schwartz, 1990). The strength of signal (SOS) corresponds to the intensity and duration of T cell activation, which is determined by multiple factors: TCR affinity, duration of antigen exposure and the availability of a co-stimulatory signal and many others (Ahmed and Nandi, 2011). SOS decides T cell fate, including proliferation, differentiation, tolerance, exhaustion, apoptosis. Previous work from our laboratory has shown that graded increase in intracellular Ca²⁺ fine-tunes activation outcomes, increases ROS and lowers cell survival (Ahmed, Mukherjee and Nandi, 2009; Ahmed et al., 2010; Joseph et al., 2024). Optimum SOS is important not only for physiological immune responses but also for therapeutic applications such as vaccinations (Butler, Nolz and Harty, 2011; Solouki *et al*., 2020) and adoptive T cell therapies (Kagoya *et al*., 2017; Zhang *et al*., 2023; Dias *et al*., 2024).

T-cell activation involves complex interaction between the TCR, APCs, costimulatory molecules, and multiple intracellular signalling pathways. Mimicking these events under physiological conditions is challenging due to the low frequency of antigen-specific T cells, the heterogeneity of T-cell populations, the complexity of T cell-APC interactions, and the diversity of signalling events that collectively determine functional outcomes (Blattman *et al*., 2002; Moon *et al*., 2007). A range of experimental models has been developed to study different aspects of T-cell activation. These include mitogenic lectins, monoclonal antibody-mediated TCR stimulation, pharmacological activators such as phorbol 12-myristate 13-acetate (PMA) and Ionomycin (P+I), antigen- and superantigen-dependent responses and TCR transgenic mice. Mice models of T cell activation are widely used as they have helped to advance our understanding of T-cell signalling, differentiation, and effector functions. The two main approaches commonly used due to their ease and cost effectiveness are: activation through αCD3 and αCD28, i.e. a TCR-dependent approach. Here, αCD3 crosslinks the TCR complex to provide primary stimulation, combined with an αCD28 antibody to deliver co-stimulatory signals that promote proliferation and cytokine production (Linsley and Ledbetter, 1993; Mukherjee *et al*., 2006). This system closely mimics physiological T cell activation by engaging the full proximal TCR signalling cascade, including Lck, ZAP-70, and LAT-dependent signalling and is widely used (Smith-Garvin, Koretzky and Jordan, 2009). The second approach uses chemical-based activation using small molecules like PMA and Ionomycin, which bypass TCR-mediated signalling. PMA is a DAG analogue that directly activates PKC thereby engaging the NF-κB and MAPK pathways. Ionomycin is a Ca^2+^ ionophore that raises intracellular Ca^2+^, thereby activating the calcineurin-NFAT axis (Smith-Garvin, Koretzky and Jordan, 2009).

Although these systems are often used interchangeably, the underlying similarities and differences in their functional outcomes remain poorly understood. Single-cell RNA sequencing has shown that human T cells stimulated using these two methods have distinct transcriptional profiles, suggesting that differences in their signalling pathways lead to changes in transcriptome profile (Lee *et al*., 2023). Comparative studies have also reported differences in cytokine profiles and proliferative output depending on the stimuli used (Olsen and Sollid, 2013; Lee *et al*., 2023). Studies in this area are important as adoptive cell therapies including chimeric antigen receptor (CAR) T cells, TCR-engineered T cells, and tumour-infiltrating lymphocyte (TIL) products all depend on the ex vivo activation and expansion of patient T cells before infusion (June *et al*., 2018; Dwivedi *et al*., 2019). Across these platforms, αCD3/αCD28 stimulation, delivered on beads or on immobilised surfaces, is the standard means of providing the two signals required to drive T cell proliferation (Trickett and Kwan, 2003; June *et al*., 2018). T cells derived from patients frequently expand sub-optimally and the resulting products may be functionally compromised or biased towards exhaustion, limiting therapeutic potency (Dwivedi *et al*., 2019; Zhang *et al*., 2023). Therefore, better understanding of intracellular signalling nodes that are rate-limiting in αCD3+αCD28-based activation is needed. It is important to address whether targeted pathway augmentation can enhance T cell expansion without promoting anergy and exhaustion. This aspect is clinically important as it applies across the spectrum of adoptive T cell therapies.

In this study, we performed a comprehensive side-by-side comparison of TCR-independent (P+I) and TCR-dependent (αCD3+αCD28) T-cell activation using primary mouse T cells. Both activation systems were evaluated at two different signal strengths and multiple aspects of T-cell responses were examined. Our results revealed that TCR-independent and TCR-dependent activation drive distinct cellular programs. Comparative analysis of these responses identified the PKC pathway as a likely point of divergence. Therefore, we explored whether selective enhancement of the PKC pathway with PMA or the Ca²⁺ pathway with Ionomycin could influence TCR-dependent activation. Understanding these effects may provide valuable insights for both basic T-cell research and the development of improved strategies for therapeutic T cell manufacturing.

## Results

### Optimisation of T cell stimulation conditions, selecting P+I concentrations and αCD3/αCD28 doses

To compare the two mechanistically distinct modes of T cell activation i.e. TCR-independent (P+I) and TCR-dependent (αCD3± αCD28), primary T cells were isolated from the lymph nodes of 6-8-week-old BALB/c mice (Figure 1A). After two rounds of panning, flow cytometry confirmed a ∼97% purity of CD3^+^ T cells (Supplementary Figure 1A), ensuring that only T cell responses were measured. The SOS is a well-established factor responsible for T cell activation, shaping cytokine production, proliferation and survival (Ahmed and Nandi, 2011). Initially, we defined optimal doses across a range of signal strengths. Earlier work from our laboratory showed that varying the Ionomycin concentration at a fixed dose of PMA produces a graded range of intracellular Ca^2+^, giving a simple, model of SOS affecting activation outcomes (Ahmed, Mukherjee and Nandi, 2009; Ahmed *et al*., 2010). In the TCR-dependent system, plate-bound αCD3 delivers a primary signal, and αCD28 co-stimulation amplifies it to give optimal activation (Linsley and Ledbetter, 1993; Mukherjee, Ahmed and Nandi, 2005; Mukherjee *et al*., 2006).

**Figure 1.**
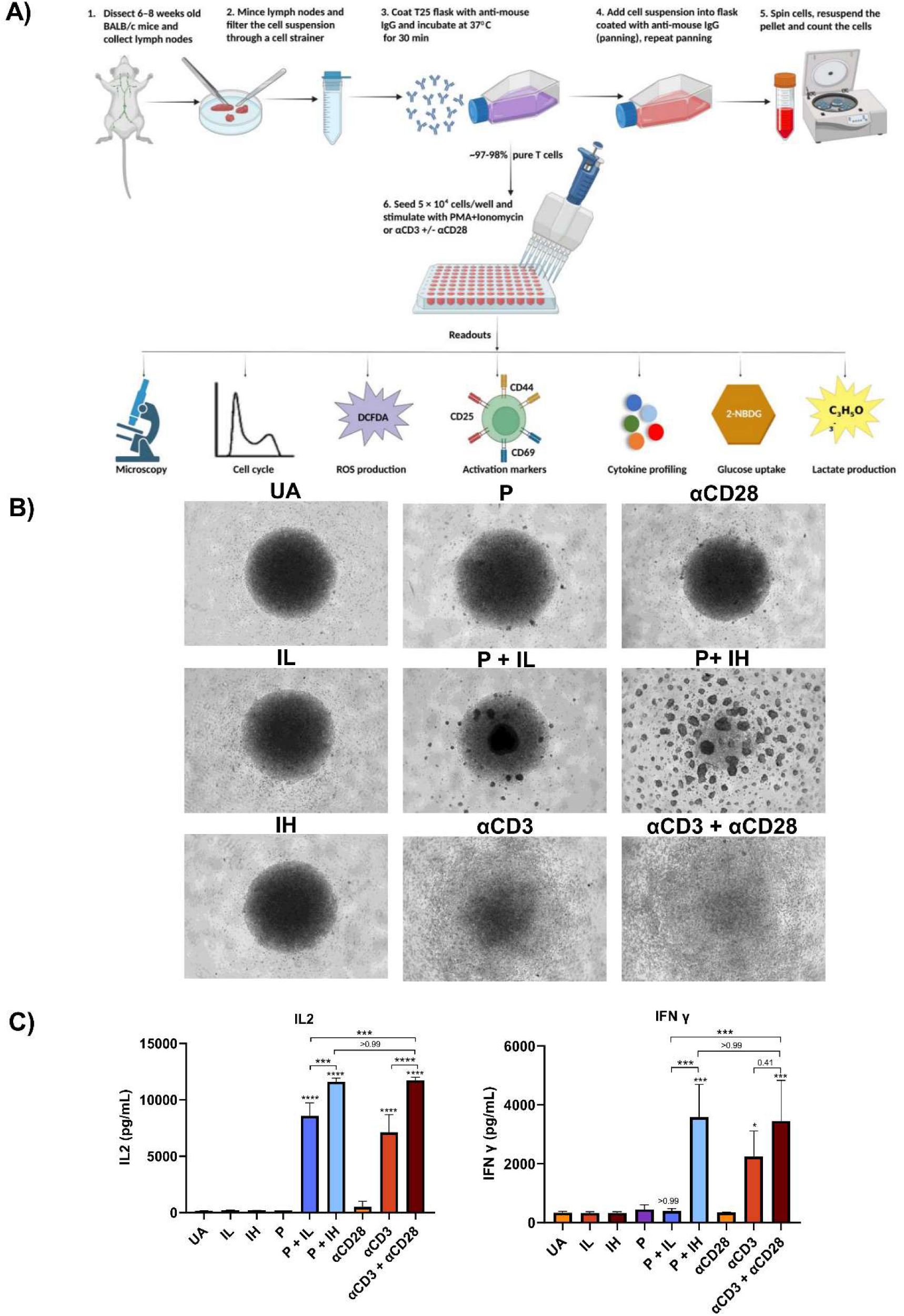
P+I and αCD3+αCD28 stimulation of T cells follows strength of signal. **(A)** Schematic representation of the experimental workflow for T cell isolation and activation using P+I or plate bound αCD3 with or without soluble αCD28 followed by downstream functional analysis. Figure 1 (A) created with BioRender.com **(B)** Representative brightfield microscopic images of T cells activated using P+I and αCD3 ± soluble αCD28 for 36 h (magnification 4×). **(C)** IL-2 and IFN-γ amounts in supernatants of activated T cells at 36 h post-activation quantified. Data are represented as mean ± SEM from three independent experiments. Statistical significance was determined by one-way ANOVA with Tukey’s multiple comparisons test. P values are multiplicity adjusted; ns p > 0.05, *p < 0.05, **p < 0.01, ***p < 0.001, ****p < 0.0001.

T cell activation-associated proliferation was measured by cell cycle analysis and expressed as the cycling-to-hypodiploidy (C:H) ratio, an index that balances productive proliferation (S + G2/M) and cell death (the sub-G1/hypodiploid fraction). We used the C:H ratio in both systems as the primary criterion for choosing conditions that maximise proliferation while limiting cell death (Joseph *et al*., 2024). For the TCR-independent system, T cells were stimulated with different PMA and/or Ionomycin combinations and scored by C:H ratio (Supplementary Figure 1B). Two conditions were chosen to represent distinct signal strengths: PMA+Ionomycin Low (P+IL), i.e. 10 ng/mL PMA with 0.2 μM Ionomycin, and PMA+Ionomycin High (P+IH), i.e. 10 ng/mL PMA with 0.8 μM Ionomycin. PMA was held constant and Ionomycin varied so that signal strength could be modulated through intracellular Ca^2+^, which has a concentration-dependent effect on activation-induced proliferation (Joseph *et al*., 2024). For the TCR-dependent system, T cells were stimulated with a range of plate-bound αCD3 concentrations, alone or with soluble αCD28, and proliferation was assessed by cell cycle analysis (Supplementary Figure 1C). Plate-bound αCD3 at 0.5 μg/mL had the highest C:H ratio for primary signal stimulation and, when combined with soluble αCD28 (1 μg/mL), amplified the initial TCR signal which led to increased proliferation and IL-2 and IFN-γ (Figure 1C).

In summary, four conditions, i.e. two systems, each at two signal strengths were established for all subsequent experiments: P+IL and P+IH for the TCR-independent system, and αCD3 alone and αCD3+ αCD28 for the TCR-dependent system. This allowed us to study and compare the two models of activation (TCR-independent versus TCR-dependent) and the SOS effects within each system to be compared directly.

## Activation induces T cell blast formation and IL-2 and IFN-γ production in an SOS-dependent manner

Post 36 hours of activation, brightfield microscopy showed clear blast formation in the P+I system in dose dependent manner (Figure 1B), confirming the dose-dependent role of Ca^2+^ in T cell activation; however, αCD3+αCD28 produced comparatively modest morphological changes, consistent with earlier reports (Joseph *et al*., 2024). Both IL-2 and IFN-γ were detectable in the supernatants at 36 hours and varied with signal strength within each system (Figure 1C). Increasing the Ionomycin concentration increased effector cytokine output (Joseph *et al*., 2024). Overall, P+IH and αCD3+αCD28 produced comparable amounts of IL-2 and IFN-γ. In the TCR-dependent system, αCD3+αCD28 induced more cytokines than αCD3 alone, consistent with the central role of αCD28 signalling in amplifying the TCR signal and driving production of various cytokines (Linsley and Ledbetter, 1993; Mukherjee, Ahmed and Nandi, 2005; Mukherjee *et al*., 2006).

### P+I stimulation induces higher expression of T cell activation markers than αCD3+αCD28

Surface expression of CD25, CD44, CD69 and PD-1 was measured across all four conditions by flow cytometry post 6 and 24 hours of activation (Figure 2, Supplementary Figure 3). CD25 is the alpha chain of IL2R, CD44 is the hyaluronic acid receptor, PD-1 is a negative regulatory receptor of T cell activation, and CD69 is the early T cell activation marker and a C-type lectin receptor. CD25 (Figure 2A, B) and CD44 (Figure 2C, D) were both upregulated in a SOS-dependent manner and were highest in the P+I system compared to αCD3+αCD28. PD-1, a T cell exhaustion marker, followed SOS and was lowest under P+IL and highest under P+IH at 36 hours; also, P+IH was comparable to αCD3+ αCD28, which correlated with strong SOS (Supplementary Figure 3A, B, C). Interestingly, CD69 behaved differently. In the P+I system its expression was not SOS-dependent; instead, high Ca^2+^ suppressed it. However, in the αCD3+αCD28 system, CD69 followed SOS and αCD28 co-stimulation significantly increased CD69 over αCD3 alone. Overall, P+IL gave the highest CD69 expression, which was significantly upregulated at 6 and 24 hours relative to the other conditions (Figure 2E, F).

**Figure 2.**
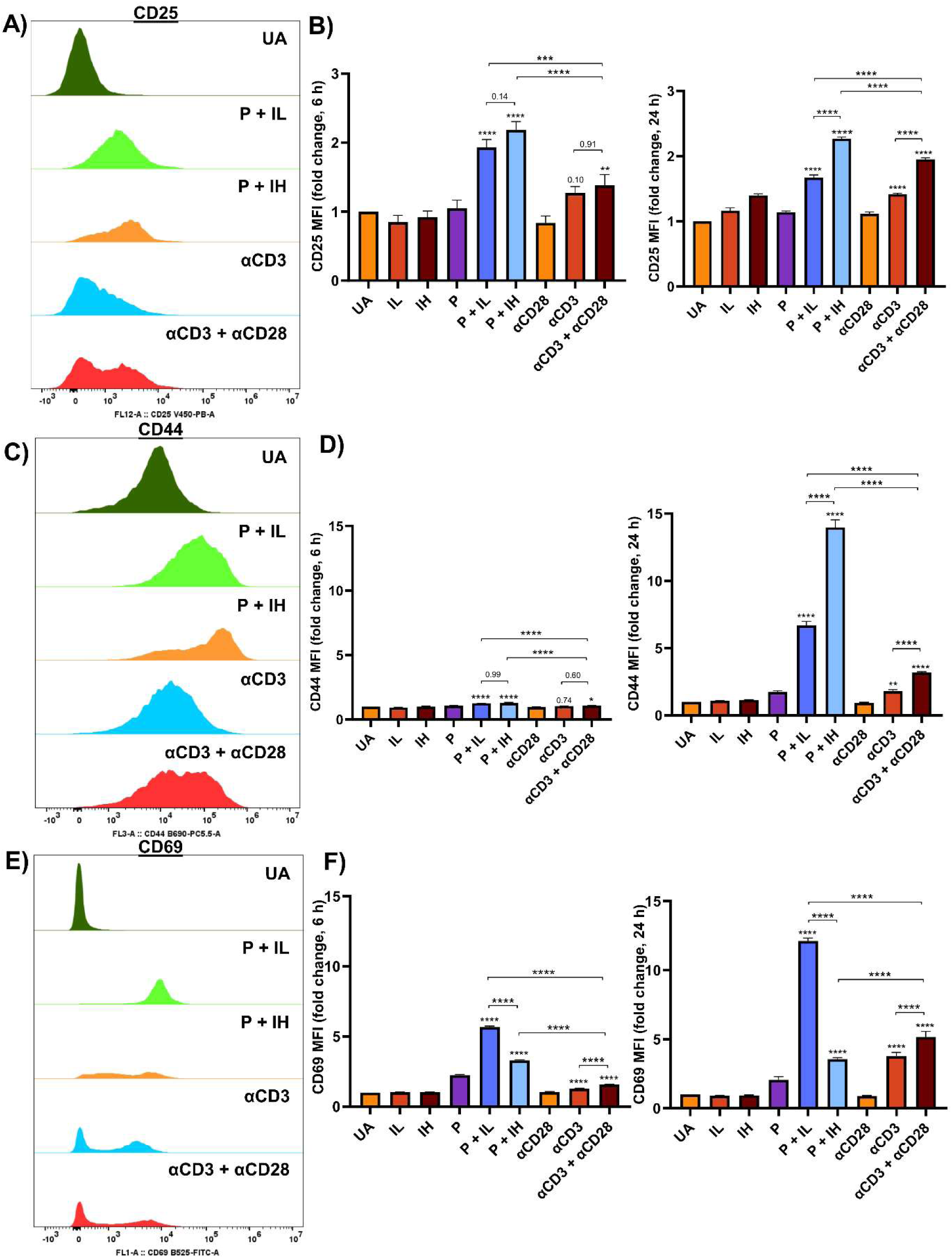
P+I stimulation leads to higher expression of T cell activation markers compared to plate-bound αCD3+αCD28 stimulation. Representative flow cytometric histograms showing surface expression of **(A)** CD25, **(C)** CD44 and **(E)** CD69 post-activation. Fold change in MFI of **(B)** CD25, **(D)** CD44, and **(F)** CD69 post-activation. All data were normalised to unactivated control. Data are represented as mean ± SEM from three independent experiments Statistical significance was determined by one-way ANOVA with Tukey’s multiple comparisons test. P values are multiplicity adjusted; ns p > 0.05, *p < 0.05, **p < 0.01, ***p < 0.001, ****p < 0.0001.

### P+I induces a greater increase in T cell size and cell cycle progression compared to αCD3+ αCD28 stimulation

To quantify differences in activation-induced proliferation, we measured cell size by forward scatter area (FSC-A) and studied the cell cycle profile using flow cytometry (Figure 3). The proportion of larger cells was significantly higher in P+I than αCD3+αCD28 at both 24 and 36 hours (Figure 3A, B). P+IL activation generated larger cells than P+IH, which is consistent with the inhibitory effects of high Ca^2+^ (Pathak *et al*., 2021; Joseph *et al*., 2024) αCD3+αCD28 stimulation resulted in a greater size increase than αCD3 alone, indicating a SOS-dependent increase in cell size. Cell cycle analysis showed that the percentage of cycling cells (S + G2/M) was significantly higher after P+I than after αCD3+αCD28 at 36 hours (Figure 3C, D). Within the P+I system, P+IL showed significantly more cycling than P+IH. Also, αCD3+αCD28 and αCD3 alone differed at 24 hours but not at 36 hours.

**Figure 3.**
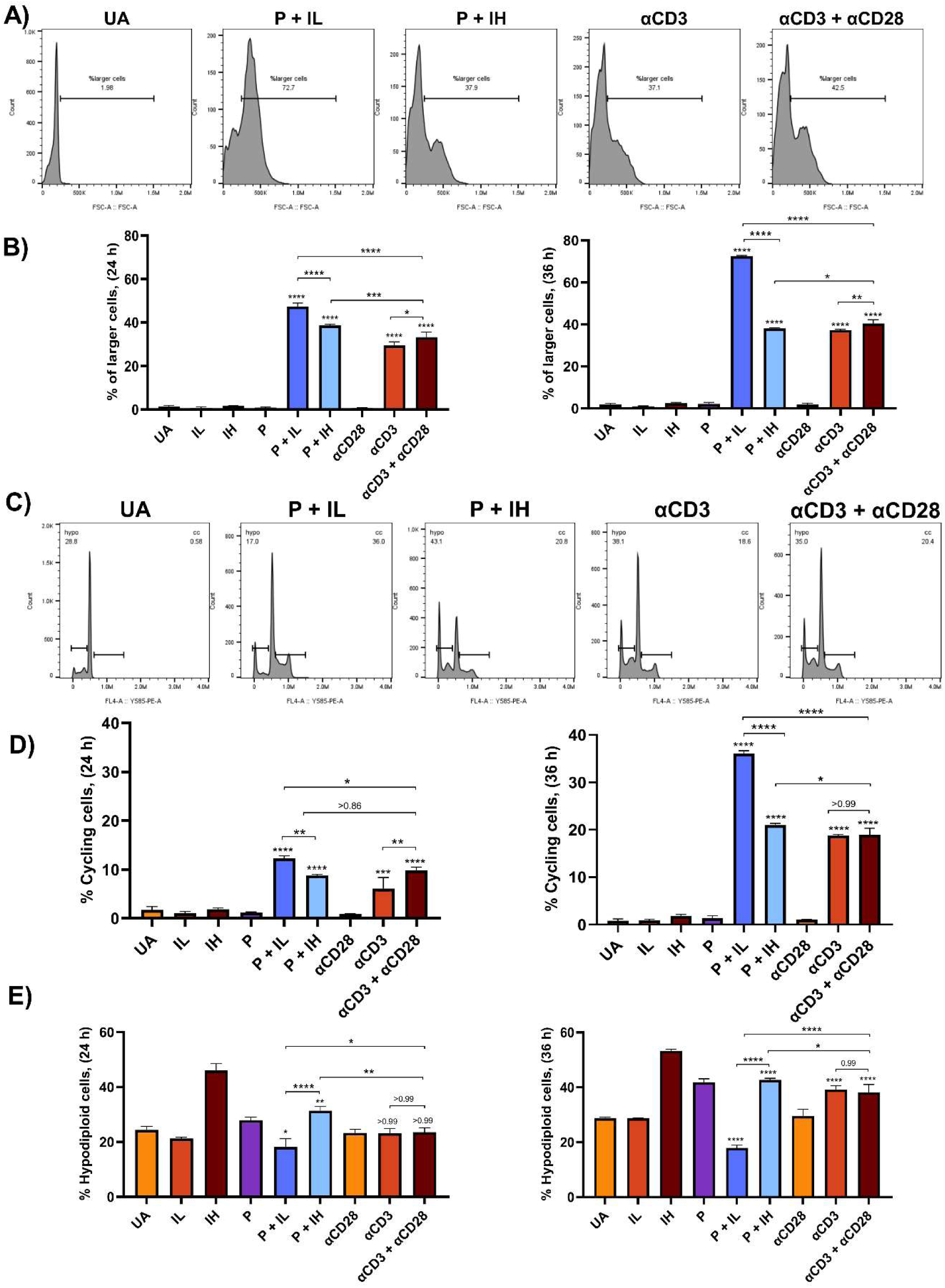
P+I activation of T cells promotes greater cell size and cell cycle progression relative to plate-bound αCD3+αCD28 stimulation. **(A)** Representative flow cytometry plots showing FSC-A distribution used to determine the proportion of larger cells compared to unactivated cells at 36 h post-activation. **(B)** The percentage of larger cells based on FSC-A parameter, representing cell size increase upon activation at indicated time points. **(C)** Representative flow cytometry plots showing cell cycle profiles at 36 h post-activation. **(D)** Percentage of cycling cells (S + G_2_M) determined by cell cycle analysis post-activation at indicated time points. **(E)** Percentage of hypodiploid (sub-G1) population derived from cell cycle analysis at indicated time points. Data are represented as mean ± SEM from three independent experiments. Statistical significance was determined by one-way ANOVA with Tukey’s multiple comparisons test. P values are multiplicity adjusted; ns p > 0.05, *p < 0.05, **p < 0.01, ***p < 0.001, ****p < 0.0001.

The hypodiploid (sub-G1) cell percentage was highest with P+IH activation (Figure 3E), consistent with our earlier finding that high intracellular Ca^2+^ raising ROS, limits proliferation and increases cell death (Joseph *et al*., 2024). Conversely, P+IL stimulation resulted in the best survival among all conditions studied. αCD3 alone and αCD3+αCD28 showed similar hypodiploidy. To measure apoptosis directly, Annexin V/PI staining was performed at 36 hours across all conditions (Supplementary Figure 2). Consistent with Figure 3E, P+IH gave the highest apoptotic fractions in this assay too, in keeping with the inhibitory effects of high intracellular Ca^2+^ (Ahmed, Mukherjee and Nandi, 2009; Joseph *et al*., 2024).

### αCD3+αCD28 stimulation shows reduced metabolic activity compared with P+I

Given these differences in cell size and cell cycle, we next assessed metabolic activity using a panel of assays (Figure 4). T cell activation is coupled to metabolic reprogramming, and signal strength decides the magnitude of this response. Glucose uptake, measured by 2-NBDG at 2 hours, was significantly higher under P+I compared to αCD3+αCD28 stimulation (Figure 4A). Within the P+I system, P+IL drove greater uptake than P+IH, reflecting both the high energy demand of activation and the dampening effects of high Ca^2+^. αCD3+αCD28 activation led to higher uptake than αCD3 alone, once again pointing to a role for αCD28 in upregulating activation (Frauwirth *et al*., 2002).

**Figure 4.**
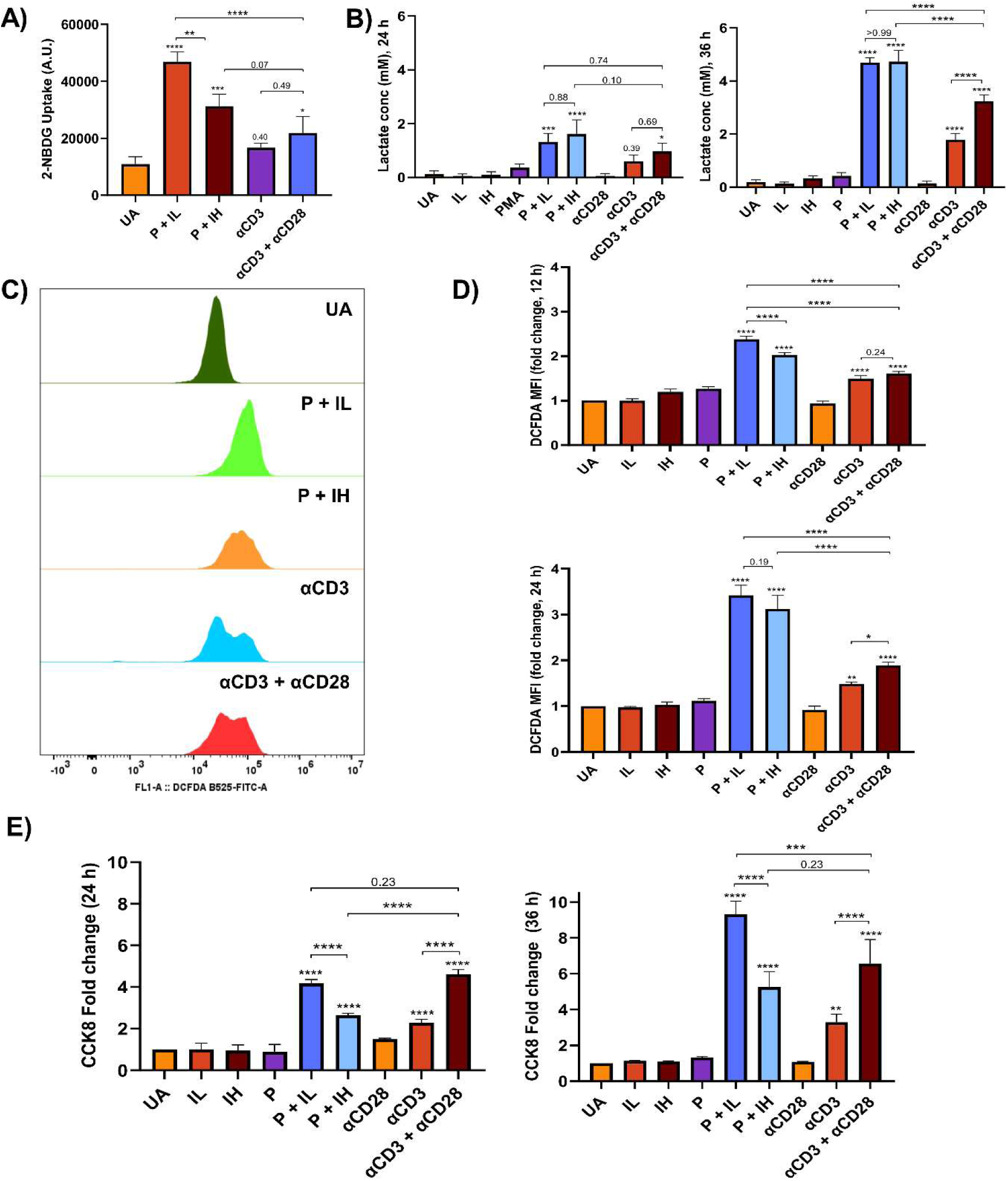
P+I activated T cells display greater metabolic activity compared to plate-bound αCD3+αCD28 stimulation. **(A)** Glucose uptake measured using 2-NBDG at 2 h post-activation. **(B)** Lactate levels in the supernatant quantified at 24 h and 36 h post-activation. **(C)** Representative flow cytometry histograms showing DCFDA fluorescence intensity at 24 h post-activation. **(D)** Intracellular ROS levels measured by DCFDA staining at 12 h and 24 h post-activation, normalised to unactivated control. **(E)** Cellular dehydrogenase activity assessed by CCK-8 assay at 24 h, and 36 h post-activation, normalised to unactivated control. Data are represented as mean ± SEM from three independent experiments. Statistical significance was determined by one-way ANOVA with Tukey’s multiple comparisons test. P values are multiplicity adjusted; ns p > 0.05, *p < 0.05, **p < 0.01, ***p < 0.001, ****p < 0.0001.

Lactate accumulation in the supernatant was significantly higher under P+I than αCD3+αCD28 at 24 and 36 hours (Figure 4B). P+IL and P+IH stimulation produced similar lactate levels, possibly due to the greater cell death with P+IH (Supplementary Figure 2). αCD3+αCD28 produced more lactate than αCD3 alone, again following SOS and consistent with a role for αCD28 in driving metabolism (Frauwirth *et al*., 2002).

Intracellular ROS, measured by 2′,7′-dichlorodihydrofluorescein diacetate (DCFDA), was significantly higher with P+I stimulation than αCD3+αCD28 at 12 and 24 hours (Figure 4C, D). P+IH produced lower ROS than P+IL at the 12-hour time point, likely reflecting dampened metabolism, in line with the glucose-uptake data. On the other hand, αCD3+αCD28 produced more ROS than αCD3 alone, again following SOS. Total cellular dehydrogenase activity (CCK-8) was higher with P+IL stimulation than P+IH at 36 hours, consistent with the above cell cycle (Figure 3D) and metabolic data (Figure 4E). αCD3+αCD28 gave an intermediate level that was significantly higher than αCD3 alone, confirming that αCD28 co-stimulation is needed for adequate metabolic activation in the TCR-dependent system (Frauwirth *et al*., 2002).

### Differential induction of cytokines and chemokine between P+I and αCD3+αCD28 stimulation

The choice of activation method strongly shapes cytokine output (Olsen and Sollid, 2013; Lee *et al*., 2023) and a panel of cytokines and chemokines was measured in 36-hour activated supernatants by ELISA (Figure 5). P+I and αCD3+αCD28 produced cytokines in an SOS- dependent manner. P+IH stimulation led to higher levels of TNF-α, IL-17 and IL-5 compared to αCD3+αCD28; however, αCD3+αCD28 produced greater levels of IL-4 and IL-10 than P+IH stimulation. IL-6 and IL-12 were detected exclusively in P+I stimulated cultures and were absent following αCD3+αCD28 stimulation. Conversely, CCL2 and IL-1β were detected only in αCD3+αCD28-stimulated cultures but not P+I. The observation that IL-6 and IL-12, but not IL-4 and IL-10, were induced greatly in P+I stimulated cultures whereas αCD3+αCD28 stimulated cultures produced IL-4 and IL-10, but not IL-6 and IL-12, clearly demonstrated that some functional responses are distinct in these two T cell activation systems.

**Figure 5.**
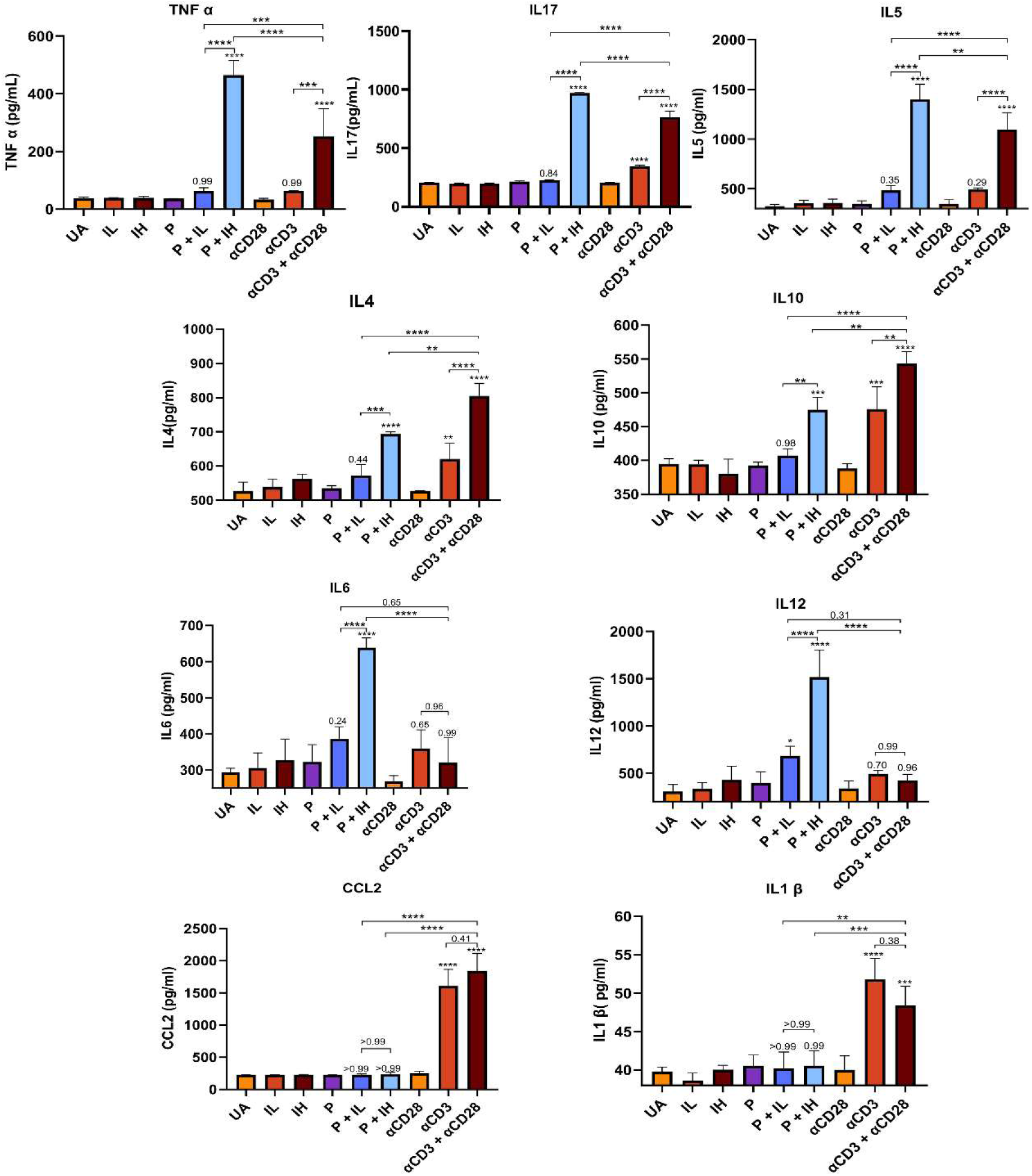
Differential expression of T cell cytokines and chemokines is observed in P+I activated cultures compared to plate-bound αCD3+αCD28 stimulation. T cells were activated using both systems for 36 h and supernatants were collected. Cytokines and chemokines in the supernatant were quantified by ELISA. Data are represented as mean ± SEM from three independent experiments Statistical significance was determined by one-way ANOVA with Tukey’s multiple comparisons test. P values are multiplicity adjusted; ns p > 0.05, *p < 0.05, **p < 0.01, ***p < 0.001, ****p < 0.0001.

### Surface CD4 expression is downregulated more strongly than CD8 upon T cell activation with P+I

CD4 and CD8 surface expression was measured by flow cytometry at 6 and 24 hours post-activation (Figure 6). Co-receptor internalisation is a hallmark of strong, PKC-driven activation and reflects the intensity of the signal (Pelchen-Matthews, Parsons and Marsh, 1993). CD4 downregulation was greatest with P+I at both time points (Figure 6A, B). αCD3+αCD28 also caused CD4 downregulation, but to a much smaller degree, and this partly recovered by 24 hours. The stronger CD4 internalisation under P+I is consistent with direct PKC activation by PMA, which bypasses the TCR requirement for receptor internalisation. CD8 downregulation (Figure 6C, D) showed mild effect with P+I stimulation, but no downregulation was seen in αCD3+αCD28. Unlike CD4, CD8 downregulation was partly reversed at the later time point.

**Figure 6.**
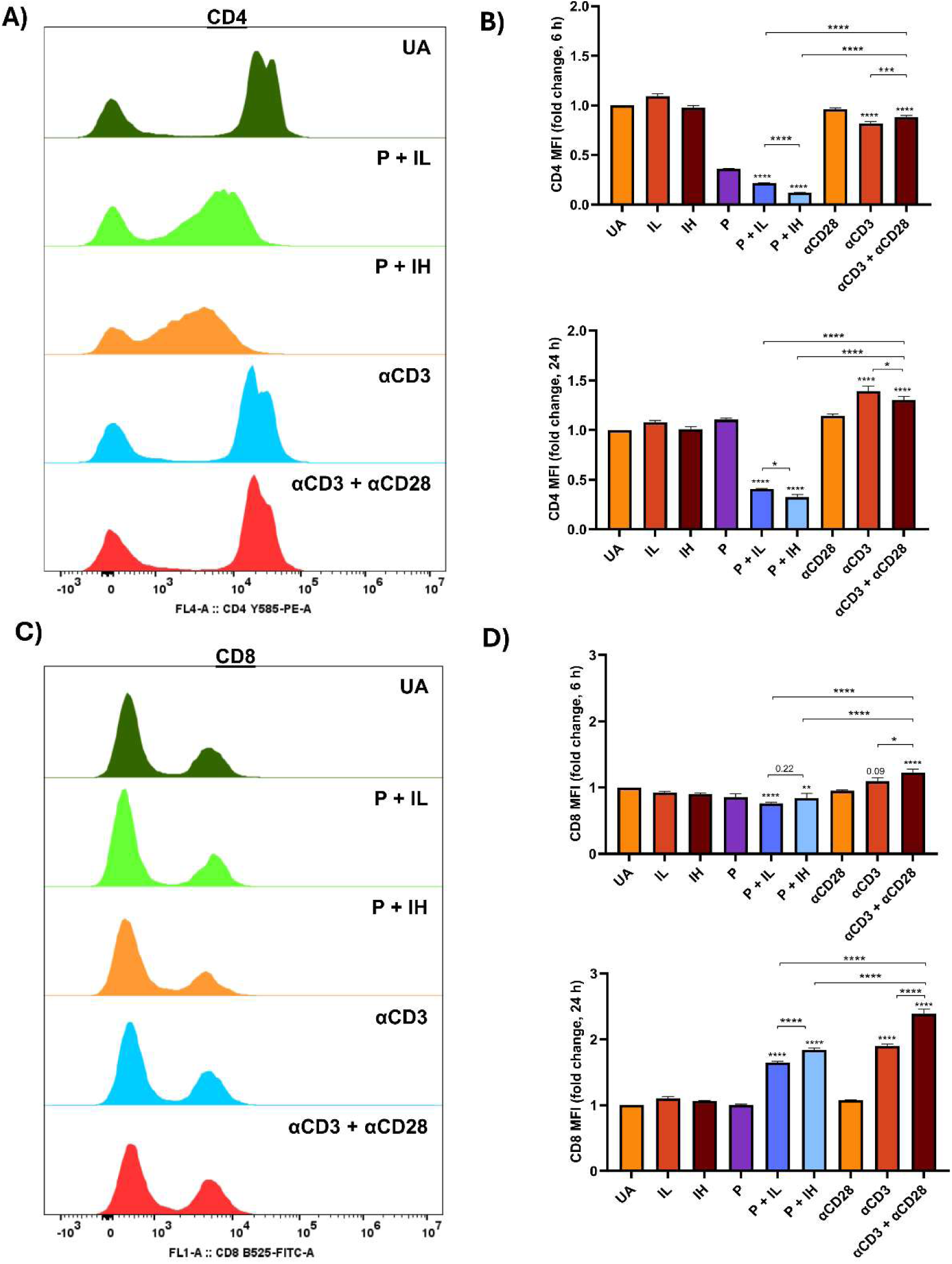
Surface expression of CD4, but not CD8, is substantially downregulated upon T cell activation with P+I. Representative flow cytometry histograms depicting **(A)** CD4 expression and **(C)** CD8 expression upon T cell activation. Fold change in MFI of **(B)** CD4 expression and **(D)** CD8. Data are represented as mean ± SEM from three independent experiments Statistical significance was determined by one-way ANOVA with Tukey’s multiple comparisons test. P values are multiplicity adjusted; ns p > 0.05, *p < 0.05, **p < 0.01, ***p < 0.001, ****p < 0.0001.

### PKC activity is significantly higher in P+I-stimulated than in αCD3+αCD28-stimulated T cells

The lower CD4 downregulation in the αCD3+αCD28 system compared to P+I stimulation prompted us to directly compare PKC activity between the two systems. PKC is the principal mediator of PMA action and main driver of the CD4 downregulation (Pelchen-Matthews, Parsons and Marsh, 1993). This observation suggested that PKC could be a key node for observed differences between the TCR-dependent and TCR-independent activation model systems. PKC activity was measured 30 minutes post-stimulation and was significantly higher with P+I stimulation compared to αCD3+αCD28 (Figure 7A, B). These data identify that a weaker PKC activation pathway may be a fundamental difference between the two systems.

**Figure 7.**
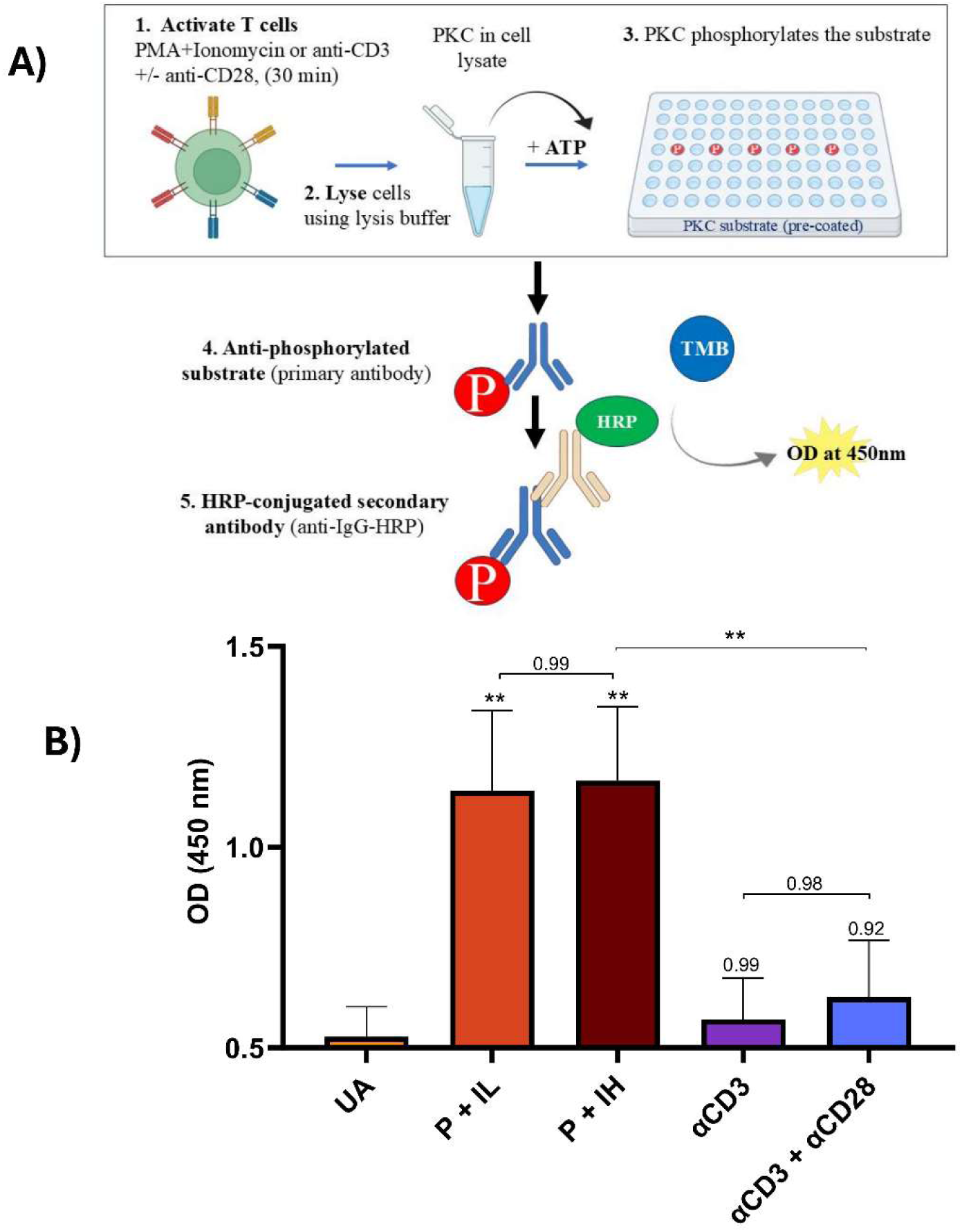
The induction of the PKC pathway is lower in plate-bound αCD3+αCD28 activation compared to P+I. **(A)** Schematic depiction of the protocol for quantification of PKC activity in T cell activation model systems. PKC activity was quantified by ELISA according to the manufacturer’s protocol. **(B)** PKC activity at 30 min post-activation. Data are represented as mean ± SEM from three independent experiments Statistical significance was determined by one-way ANOVA with Tukey’s multiple comparisons test. P values are multiplicity adjusted; ns p > 0.05, *p < 0.05, **p < 0.01, ***p < 0.001, ****p < 0.0001

### Supplementing the αCD3+αCD28 system with PMA increases proliferation, metabolic activity and surface activation markers

To test directly whether boosting PKC signalling rescued the weaker responses in the αCD3+ αCD28 system, T cells were supplemented with exogenous PMA (1, 10 or 50 ng/mL) or Ionomycin (0.2, 0.4 or 0.8 μM), and surface expression markers, cell cycle progression and metabolic activity were assessed (Figure 8, 9, 10). CD25, CD44 and CD69 were measured after 24 hours in αCD3+ αCD28-stimulated T cells treated with PMA or Ionomycin (Figure 8). PMA in a dose-dependent manner significantly increased CD25 (Figure 8A), CD44 (Figure 8B) and CD69 (Figure 8C), showing that increasing PKC signalling in the TCR-dependent system increased activation-marker expression. Ionomycin did not significantly rescue CD25 or CD44 at any concentration tested; however, it reduced CD69 expression in dose dependent manner, consistent with the inhibitory effects of high intracellular Ca^2+^ (Pathak *et al*., 2021; Joseph *et al*., 2024). Surprisingly, PMA supplementation decreased PD-1 expression (Supplementary Figure 3A, B).

**Figure 8.**
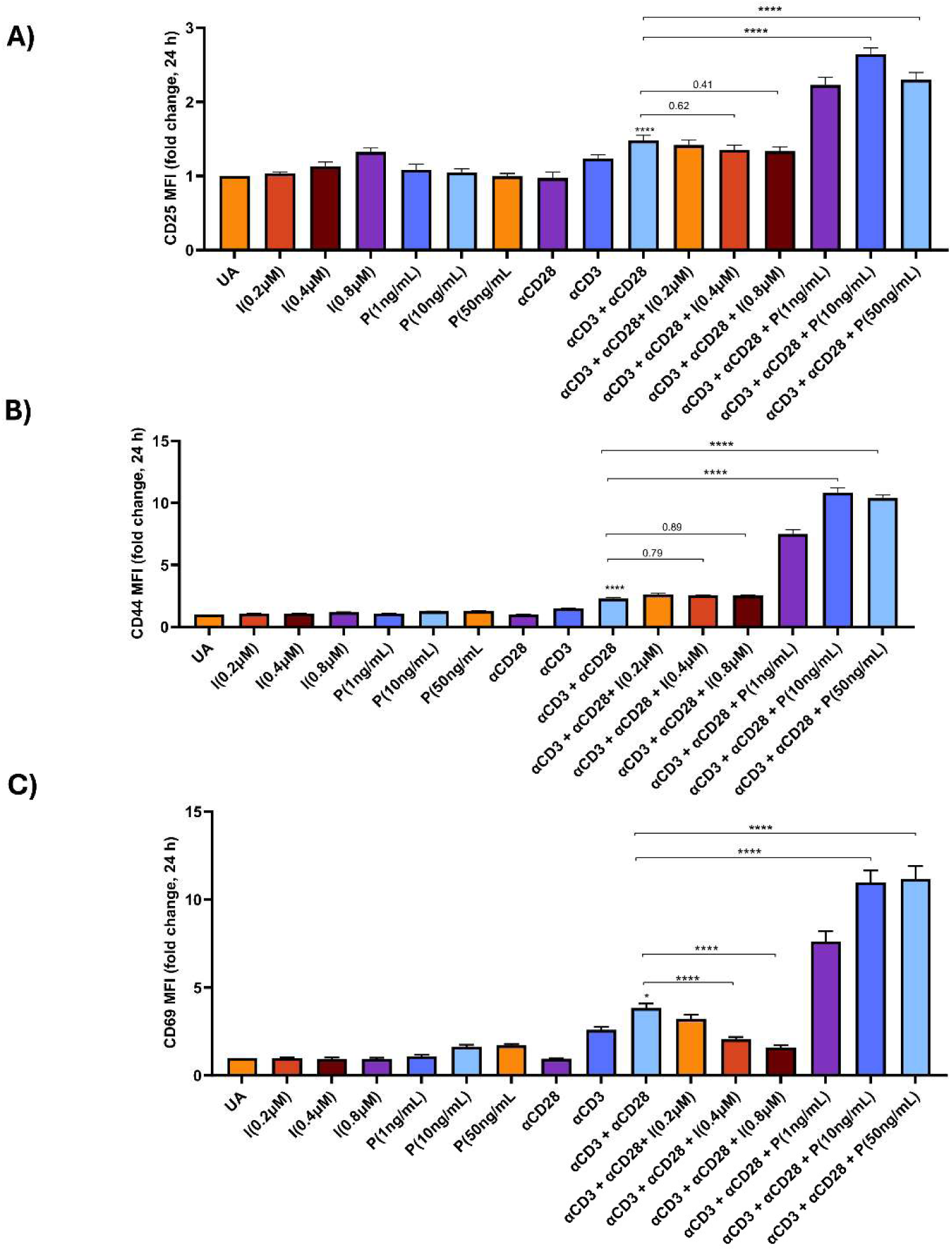
Addition of PMA, but not Ionomycin, enhances surface activation markers in T cells activated with αCD3+αCD28. Fold change in MFI of **(A)** CD25, **(B)** CD44, and **(C)** CD69 expression at 24 h post-activation. All data are normalised to unactivated control. Data are represented as mean ± SEM from three independent experiments Statistical significance was determined by one-way ANOVA with Tukey’s multiple comparisons test. P values are multiplicity adjusted; ns p > 0.05, *p < 0.05, **p < 0.01, ***p < 0.001, ****p < 0.0001.

**Figure 9.**
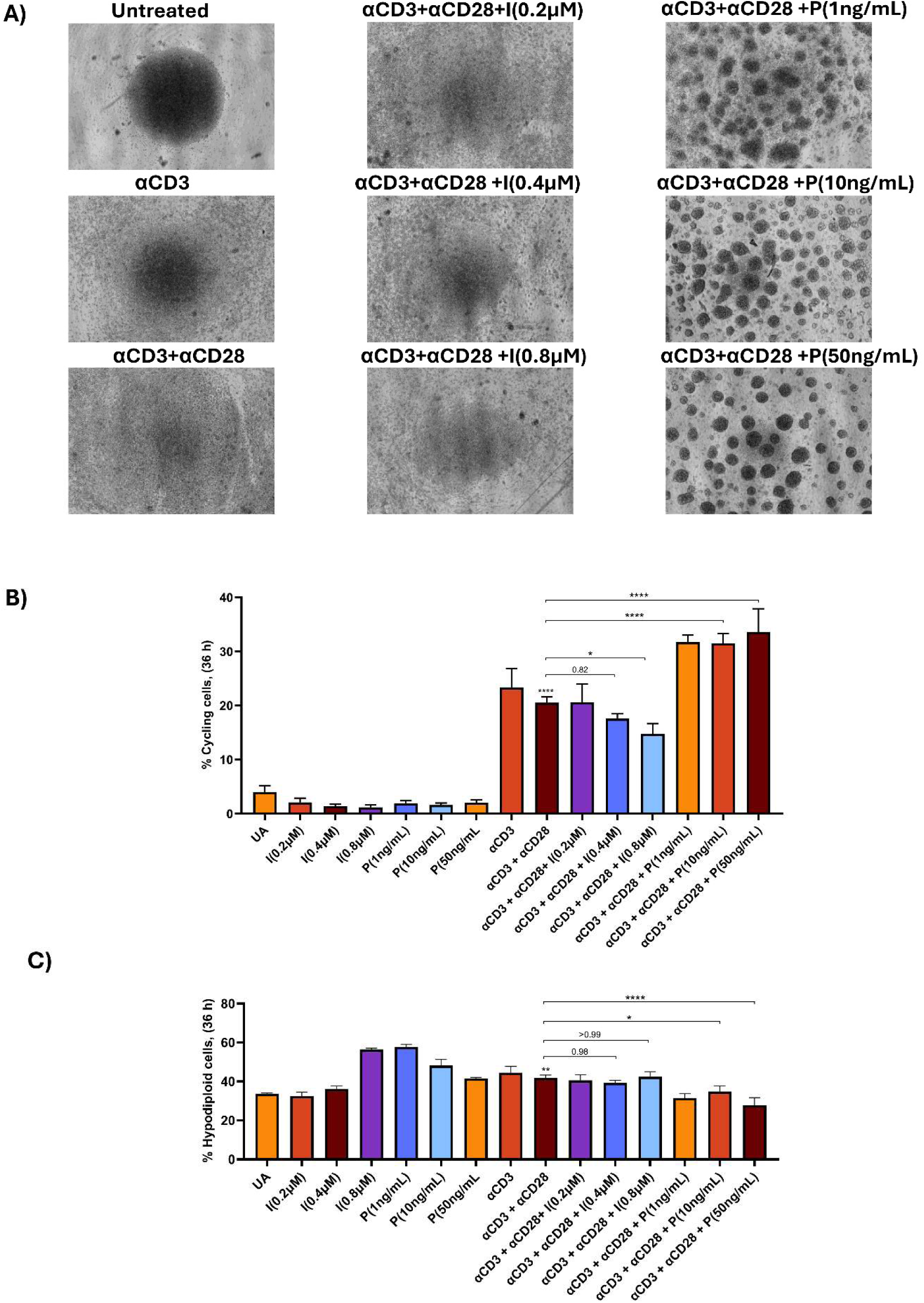
Addition of PMA increases T cell cycling upon plate-bound αCD3 + αCD28 activation. **(A)** Representative brightfield microscopic images of T cells activated with plate-bound αCD3 + αCD28 and supplemented with indicated concentrations of PMA and Ionomycin at 36 h post-activation (magnification 4×). **(B)** Percentage of cycling cells (S + G_2_M) determined by cell cycle analysis at 36 h post-activation. **(C)** Percentage of hypodiploid (sub-G1) population derived from cell cycle analysis. Data are represented as mean ± SEM from three independent experiments Statistical significance was determined by one-way ANOVA with Tukey’s multiple comparisons test. P values are multiplicity adjusted; ns p > 0.05, *p < 0.05, **p < 0.01, ***p < 0.001, ****p < 0.0001.

Addition of PMA to αCD3+αCD28 stimulated cultures also increased blast formation and the proportion of cycling cells (S + G2/M) at 36 hours (Figure 9A, B) and reduced hypodiploidy (Figure 9C). On the other hand, Ionomycin did not increase cycling, but at 0.8 μM increased hypodiploidy reproducing the inhibitory effects of high intracellular Ca^2+^ (Ahmed, Mukherjee and Nandi, 2009; Joseph *et al*., 2024).

Also, addition of PMA significantly increased lactate production, dehydrogenase activity and intracellular ROS in the αCD3+αCD28 system (Figure 10A, B, C). Conversely, Ionomycin did not improve metabolism, but rather reduced dehydrogenase activity, which is consistent with the inhibitory effects of high intracellular Ca^2+^ (Ahmed, Mukherjee and Nandi, 2009; Joseph *et al*., 2024). Together, these data show that insufficient PKC signalling is the primary deficit of the αCD3+αCD28 system relative to P+I.

**Figure 10.**
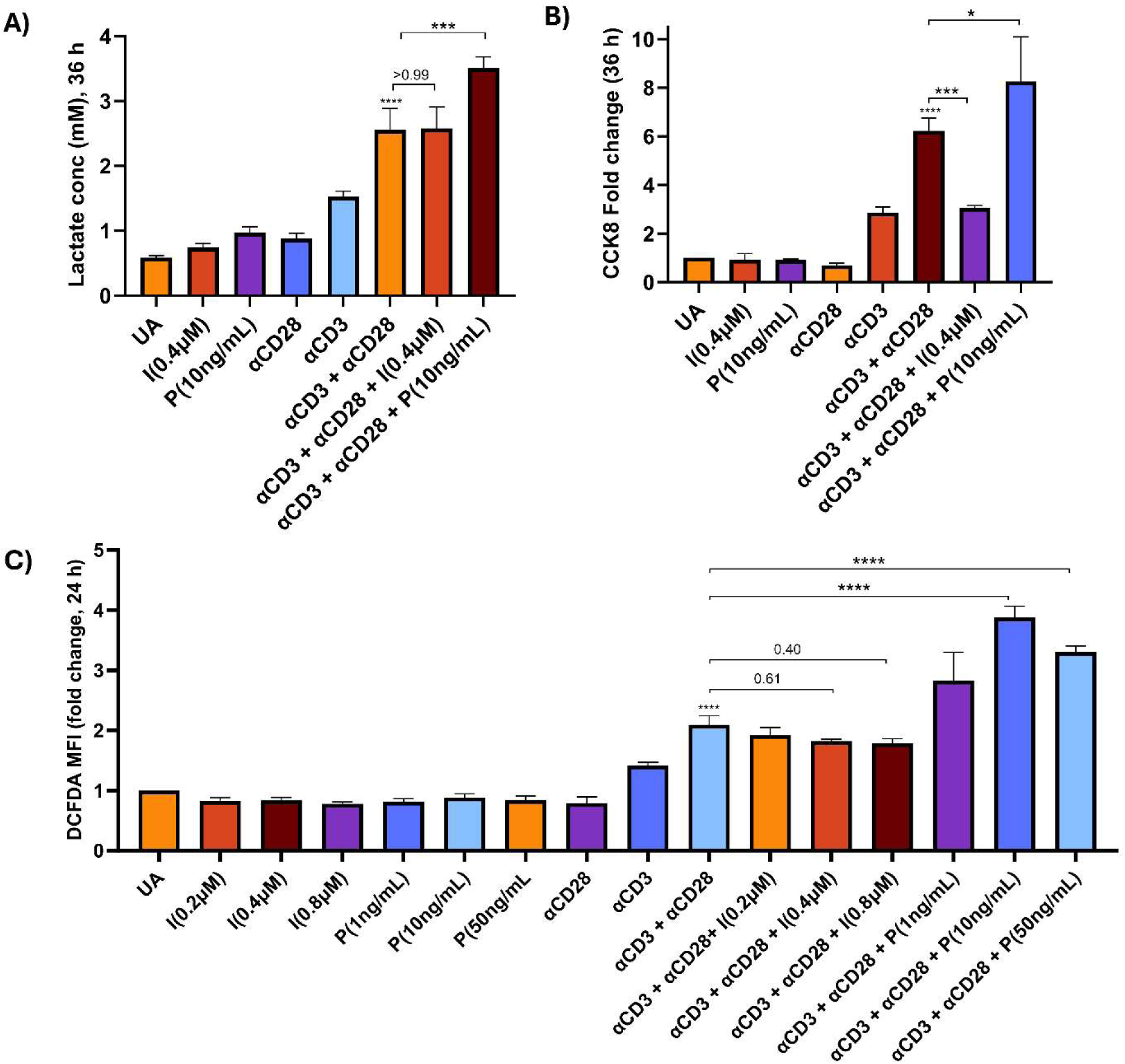
Addition of PMA increases metabolic activity in T cells activated with αCD3 + αCD28. **(A)** Lactate amounts in the supernatant post 36 h of activation in the αCD3 + αCD28 system supplemented with PMA or Ionomycin. **(B)** Cellular dehydrogenase activity assessed by CCK-8 assay at 36 h post-activation. **(C)** Intracellular ROS levels measured by DCFDA staining at 24 h post-activation. All data are normalised to unactivated control. Data are represented as mean ± SEM from three independent experiments Statistical significance was determined by one-way ANOVA with Tukey’s multiple comparisons test. P values are multiplicity adjusted; ns p > 0.05, *p < 0.05, **p < 0.01, ***p < 0.001, ****p < 0.0001.

### PMA or Ionomycin supplementation differentially modulates cytokine profiles in the αCD3+ αCD28 system

Cytokine and chemokine production were measured post 36-hour of stimulation of T cells with αCD3+αCD28 together with PMA or Ionomycin (Figure 11). PMA increased TNF-α, IL-4, IL-5 and IL-10 while decreasing IL-17, CCL2 and IL-1β. On the other hand, Ionomycin decreased IL-2, TNF-α, IL-5, CCL2 and IL-1β. Together, these data clearly demonstrate that differential effects of PMA and Ionomycin in T cells stimulated with αCD3+αCD28. It is likely that PKC signalling is sub-optimal in αCD3+αCD28 stimulated cultures and enhancing PKC signalling increases T cell activation.

**Figure 11.**
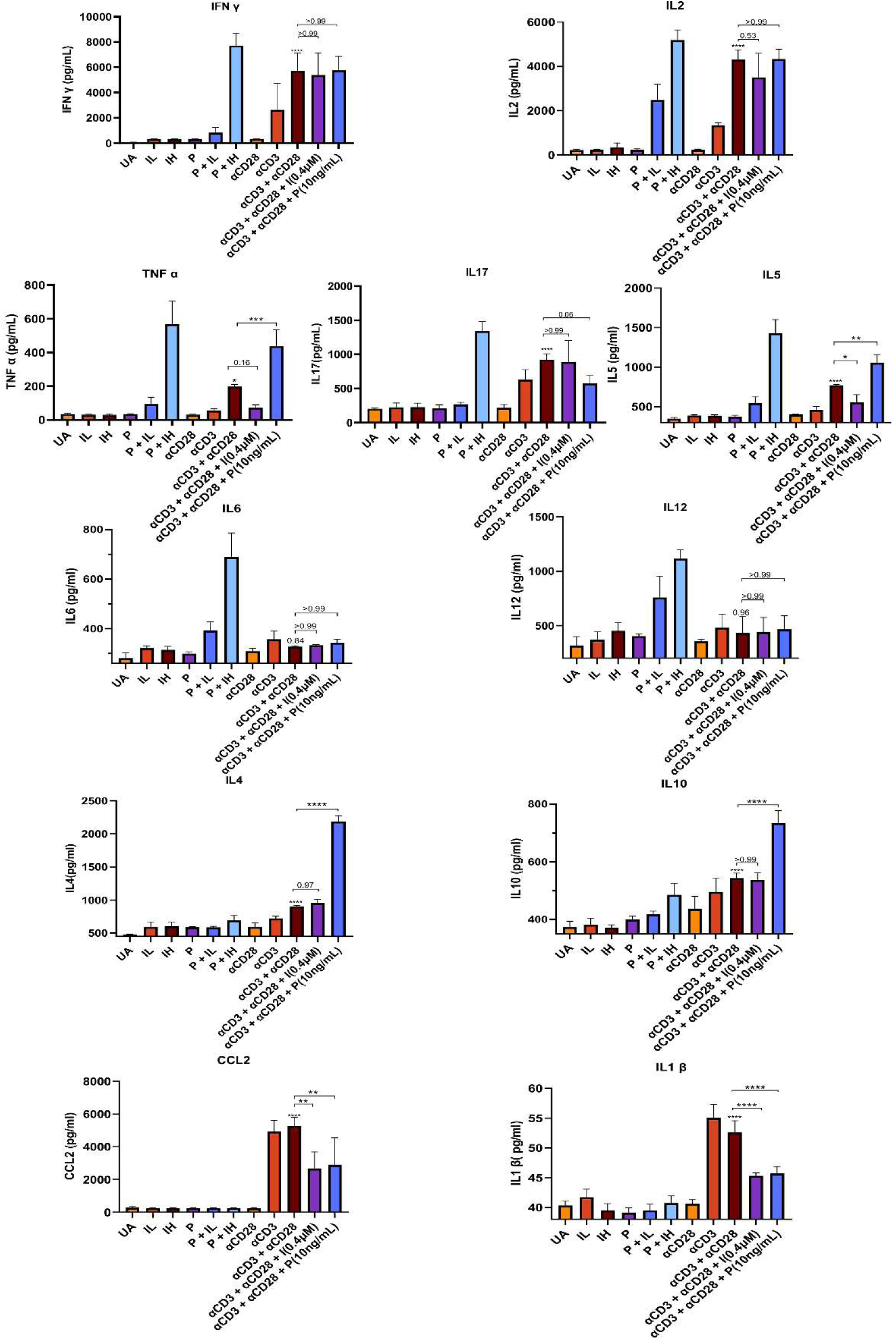
Addition of PMA or Ionomycin differentially modulates the induction of cytokines and chemokines in T cells activated with αCD3+αCD28. T cells were activated using the αCD3+αCD28 system supplemented with PMA or Ionomycin for 36 h and supernatants were collected. Cytokines and chemokines in the supernatant were quantified by ELISA. Data are represented as mean ± SEM from three independent experiments Statistical significance was determined by one-way ANOVA with Tukey’s multiple comparisons test. P values are multiplicity adjusted; ns p > 0.05, *p < 0.05, **p < 0.01, ***p < 0.001, ****p < 0.0001.

## Discussion

In this study we compared the TCR-dependent (αCD3+αCD28) and TCR-independent (P+I) T cell activation responses under identical conditions. A comprehensive comparative analysis across a range of signal strengths in these T cell culture systems demonstrate that T cells show consistent and reproducible differences in cell morphology, cell size, cell cycle progression, expression of surface activation markers, metabolism, apoptosis and cytokine profiles. P+I mediated activation produced quantitatively stronger responses than αCD3+αCD28 mediated activation across most readouts (Figures 2-6). What is perhaps more informative than the magnitude of this difference is its mechanistic basis: when selectively supplemented with PMA, but not Ionomycin, the αCD3+αCD28 system showed remarkable restoration of the proliferative, metabolic and phenotypic deficits, highlighting the PKC signalling arm as the rate-limiting step in αCD3+αCD28 dependent T-cell activation (Figure 12).

**Figure 12.**
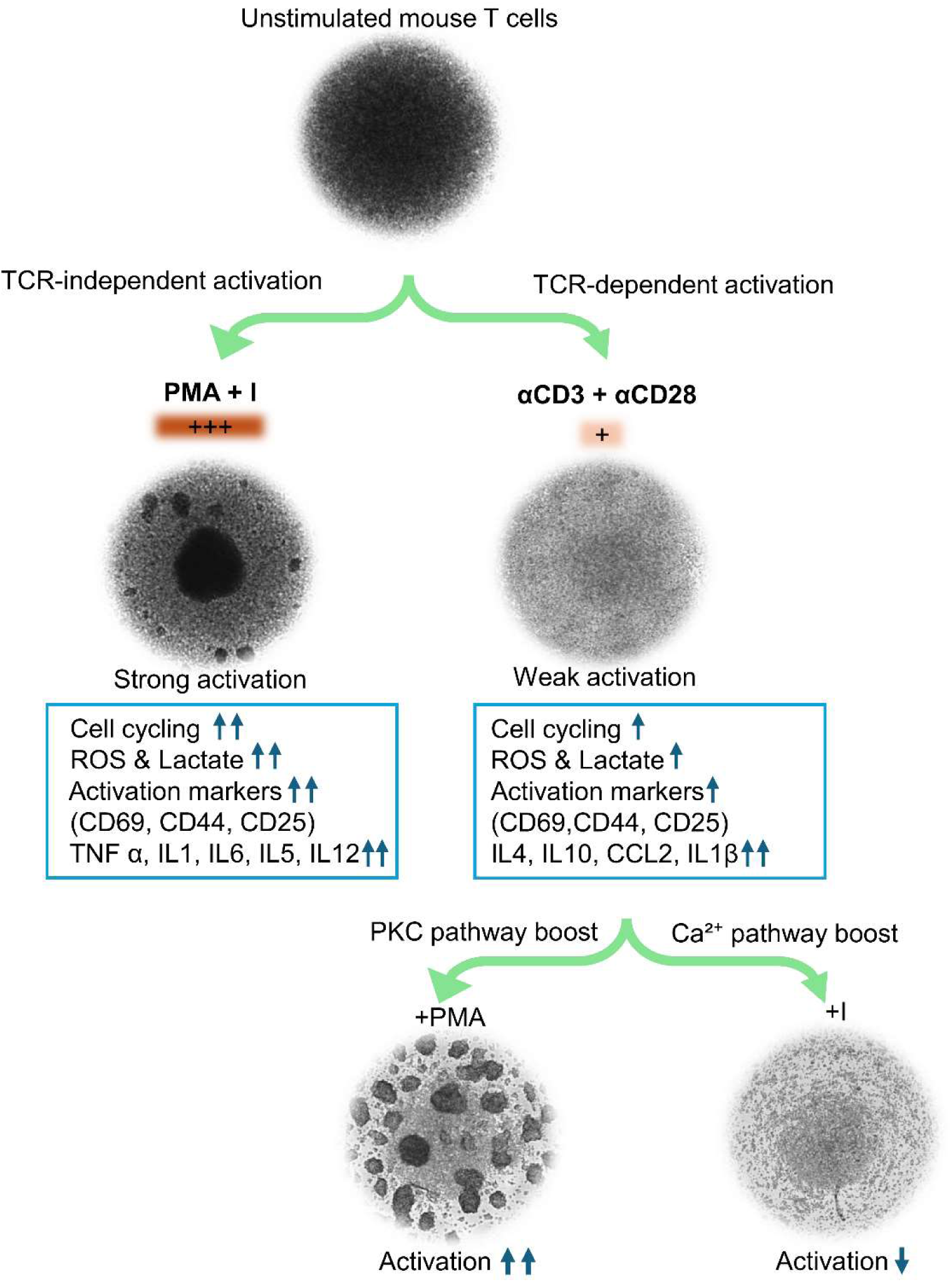
Graphical illustration summarising the comparative analysis of the two T cell activation systems: TCR-dependent (αCD3+αCD28) and TCR-independent (PMA+Ionomycin). This study highlights the identification of PKC as a critical pathway that enhances TCR dependent activation via αCD3 + αCD28.

Although optimum concentrations chosen by C:H ratio produced comparable IL-2 and IFN-γ levels in both systems following SOS, other parameters such as blast formation showed remarkable differences, with P+I activation system outperforming the αCD3+αCD28 system (Figure 1). The αCD3+αCD28 system followed SOS and exceeded αCD3 activation alone in terms of cell size (Figure 1), total dehydrogenase activity, lactate production, and IL-2, IL-4, IL-5, IL-10, IL-17 and TNFα production (Linsley and Ledbetter, 1993; Mukherjee *et al*., 2006). In contrast, P+I system did not follow SOS: P+IH showed lower cell size and cycling compared to P+IL, along with the highest hypodiploidy, demonstrating that high Ca^2+^ inhibits rather than enhances these parameters (Figure 3, Figure 4, Figure 5). This confirms the concentration-dependent effects of intracellular Ca²⁺, where moderate levels support proliferation while higher levels inhibit cell cycle progression and promote apoptosis (Ahmed, Mukherjee and Nandi, 2009; Joseph *et al*., 2024).

Cell surface markers are known to be modulated upon T cell activation. CD25, CD44 and PD-1 followed SOS in both systems and were highest under P+IH compared with αCD3+αCD28. Notably, CD44 surface upregulation was far greater under P+I compared to αCD3+αCD28 at the later time point (Figure 2). PD-1, a well-established marker of T cell exhaustion (Wherry, 2011; Wherry and Kurachi, 2015), was notably lower in P+IL, the condition that also showed optimal proliferation. It is possible that an appropriate signal strength can drive robust activation while limiting the induction of exhaustion markers. In contrast, αCD3+αCD28 showed PD-1 levels comparable to P+IH, raising the possibility that standard TCR-based stimulation may drive T cells towards exhaustion (Supplementary Figure 3). In addition, we found that PMA supplementation increased CD25, CD44 and CD69 expression but not PD-1, which suggests that the cells were proliferating and activating without moving towards exhaustion which may have useful implications for improving T cell responses in therapeutic settings (June *et al*., 2018). Interestingly, CD69 behaved distinctly from other markers: while it followed SOS within the αCD3+αCD28 system, it was suppressed with P+IH activation in the TCR-independent cell culture system (Figure 2). This pattern was also mirrored in supplementation studies where Ionomycin add-back similarly reduced CD69 surface expression (Figure 8). It appears that CD69 expression closely paralleled the metabolic activity pattern across all conditions, consistent with its emerging role not merely as an early activation marker but as a metabolic gatekeeper regulating cellular metabolism (Cibrian and Sanchez-Madrid, 2017). The mechanism by which high Ca^2+^ inhibits CD69 expression remains unknown and represents a novel observation of the present study.

T cell activation is metabolically demanding. Upon activation, T cells shift from oxidative phosphorylation toward aerobic glycolysis, called the Warburg effect. This increases glucose uptake, lactate production, ROS generation and mitochondrial activity to meet the biosynthetic requirements of rapid proliferation. Studies have shown that T cell activation induced ROS is PKC theta dependent (Kamiński *et al*., 2007). This metabolic program scales with signal strength via CD28-PI3K-AKT-mTOR signalling pathway (Frauwirth *et al*., 2002). Thus, activation methods that engage PI3K to different extends are likely to produce distinct metabolic states. These different activation systems may lead to differential functional, phenotypic and metabolic outcomes, highlighting the need for direct comparison (Chang *et al*., 2013; Buck, O’Sullivan and Pearce, 2015; O’Neill, Kishton and Rathmell, 2016). In line with the stronger proliferative response, P+I showed greater metabolic activity than αCD3+αCD28 system across multiple readouts: glucose uptake, lactate accumulation, intracellular ROS and total dehydrogenase activity (Figure 4). αCD3+αCD28 exceeded αCD3 alone across all these parameters, consistent with the established role of CD28 signalling in upregulating glucose transporter expression and driving glycolytic flux (Frauwirth *et al*., 2002). However, P+IH failed to enhance metabolic activity significantly over P+IL, consistent with prior reports from our group showing that high intracellular Ca²⁺ dampens T cell activation (Ahmed, Mukherjee and Nandi, 2009; Joseph *et al*., 2024).

The two activation systems generated distinct cytokine and chemokine profiles, further demonstrating that TCR-dependent and TCR-independent activation are not functionally equivalent (Figure 5). Compared with αCD3+αCD28, P+IH induced higher production of the inflammatory cytokines TNF-α, IL-17 and IL-5, whereas αCD3+αCD28 favoured a more balanced cytokine profile with greater IL-4 and IL-10 production which is consistent with previous studies (Olsen and Sollid, 2013; Lee *et al*., 2023). The higher IL-4 levels are consistent with the established role of CD28 co-stimulation in promoting IL-4 expression through the GATA-3 pathway (Rulifson *et al*., 1997), while the lower IL-10 observed following P+I agrees with reports that high doses of PMA/Ionomycin stimulation suppresses IL-10 production (Olsen and Sollid, 2013). Importantly, supplementation of the αCD3+αCD28 system with PMA selectively enhanced the production of TNF-α, IL-5, IL-4 and IL-10 (Figure 11), closely paralleling the restoration of proliferation, metabolic activity and activation-marker expression observed under the same conditions, further supporting the conclusion that limited PKC signalling is a major factor restricting the functional response following αCD3+αCD28 stimulation. IL-6 expression in activated T cells is driven by PKC-θ-dependent activation of NF-κB (Sofi *et al*., 2009; Vemulawada *et al*., 2026). Exclusive production of IL-6 in P+I stimulation therefore aligns with the greater PKC activity generated by this system in comparison to TCR-dependent αCD3+αCD28 stimulation. Although helper T cells, NKT cells and γδ T cells are recognised secondary sources of IL-12, its dominant producers are myeloid dendritic cells and macrophages, which generate IL-12 through NF-κB-driven transcription (Macatonia *et al*., 1995; Trinchieri, 2003). The exclusive production of IL-12 under P+I is most likely explained by TCR independent pharmacological activation. T-cell-derived CCL2 has previously been reported in tumour-associated and chronic inflammatory settings (Owen *et al*., 2005, 2011), and its production has largely been attributed to disease-specific microenvironmental cues. Another study similarly identified T cells as a source of CCL2 within the tumour microenvironment only in the context of pathological stimulation (Jin *et al*., 2021). To our knowledge, the present study is the first to demonstrate CCL2 production directly following physiological αCD3+αCD28 stimulation of healthy primary T cells under defined *in vitro* conditions, independent of tumour- or disease-associated factors. Notably, CCL2 was completely absent following P+I stimulation. Importantly, supplementation with either PMA or Ionomycin inhibited its production, indicating that these compounds suppress the TCR-dependent CCL2 production by T cells. IL-1β is an innate immune cytokine produced by myeloid cells; however, recent studies indicate that activated T cells can also express it under specific conditions. Mechanistically, TCR stimulation has been shown to induce pro-IL-1β expression in Th17 cells, while subsequent inflammasome activation promotes its maturation and release through a T-cell-intrinsic ASC–NLRP3–caspase-8 pathway (Martin *et al*., 2016). Consistent with this, we detected low but measurable levels of IL-1β exclusively following αCD3+αCD28 stimulation, whereas none was detected after P+I treatment, suggesting that TCR/CD28 signalling may preferentially engage pathways leading to IL-1β induction. This raises the possibility that T cells themselves contribute to IL-1β production following physiological activation, although the underlying signalling mechanisms require further investigation.

Activation-induced co-receptor modulation differed greatly between the two activation systems and provides an independent phenotypic readout of differential PKC signalling. CD4 was downregulated far more extensively following P+I than αCD3+αCD28 stimulation, consistent with the established PKC-dependent pathway in which PKC phosphorylation promotes CD4 dissociation from p56lck, clathrin-mediated internalisation and subsequent lysosomal degradation (Pelchen-Matthews *et al*., 1992; Pelchen-Matthews, Parsons and Marsh, 1993; Pitcher *et al*., 1999). As this entire endocytic cascade is driven by PKC-mediated phosphorylation of the CD4 cytoplasmic tail, the much greater CD4 loss in the P+I system is consistent with the stronger, sustained PKC activation generated by direct phorbol ester stimulation, relative to the weaker, receptor-limited PKC engagement in the TCR-dependent pathway (Figure 6). The recovery kinetics further distinguished the two activation models: CD4 expression returned to near-baseline levels after αCD3+αCD28 stimulation but recovered only partially after P+I (Figure 6). This pattern is consistent with a model in which transient, regulated PKC signalling supports reversible CD4 internalisation with recycling, whereas the combination of sustained PKC activation Ca²⁺ engages a more durable downregulation pathway, resulting in sustained CD4 loss (Anderson and Coleclough, 1993). These findings suggest that αCD3+αCD28 primarily induces reversible co-receptor internalisation, whereas P+I promotes more persistent CD4 loss. Sustained CD4 downregulation is a documented *in vivo* outcome of chronic antigen exposure with real biological consequences, establishing a negative-feedback context in which the strongest stimulus drives the greatest co-receptor loss yet the least productive proliferative outcome (Beaumier *et al*., 2009; Grishkan *et al*., 2013). On the other hand, CD8 regulation followed a distinct pattern compared to CD4: first downregulation of CD8, compared to CD4, was minimal and observed only in P+I at early time point. Second, CD8 surface expression was increased at the later time point with P+I and αCD3+αCD28 (Figure 6). Together, these findings indicate that CD4 is readily regulated by PKC-mediated internalisation, whereas CD8 downregulation requires the combined effects of strong PKC activation and Ca²⁺ signalling, reflecting their differential mechanistic wiring.

Collectively, these findings demonstrate that P+I and αCD3+αCD28 are distinct models of T-cell activation. In line with previous reports (Olsen and Sollid, 2013; Lee *et al*., 2023) our results show that both the type and strength of the activation signal shape the cellular response, and the choice of stimulation method and its intensity should therefore be guided by the question being addressed. As the two systems generate distinct signalling, functional and metabolic states, results obtained with one model may not necessarily apply to the other, a consideration particularly relevant when studying pharmacological modulators of T-cell activation, where the stimulation method can influence both the efficacy and mechanism of action of a compound. The findings also carry direct implications for adoptive T-cell therapies, particularly CAR-T cell manufacturing. For T cell therapies, TCR-dependent activation via plate-bound or bead-conjugated anti-CD3/anti-CD28 is the conventional method for T cell expansion (Trickett and Kwan, 2003; June *et al*., 2018). Despite its widespread clinical use, this system has its own challenges: patient-derived T cells often expand poorly and face different challenges such as low numbers and exhaustion (June *et al*., 2018; Zhang *et al*., 2023; Mehta *et al*., 2024). Our data suggest that selectively enhancing the PKC arm with PMA rescues these deficits without causing exhaustion thus identifying PKC signalling as a rational and targetable arm for improving TCR-dependent activation. This enhancement may increase the transfection efficiency and also reduce the therapy time. However, PMA is a tumor promoter and there is a need to consider other compounds that enhance PKC signalling (Le *et al*., 2009; Wang *et al*., 2026). These observations raise the testable hypothesis that controlled reinforcement of the PKC arm during the activation phase of CAR-T manufacturing, using clinically acceptable PKC agonists, could improve T-cell expansion.

Some limitations of this study need to be acknowledged. Experiments were conducted with primary mouse T cells from BALB/c mice, comprising both CD4⁺/CD8⁺ naïve and effector populations and the relative contributions of different subsets to the observed functional and metabolic differences were not resolved. In future, subset-specific analyses may reveal additional nuances. Also, we need to screen and identify additional cytokine, chemokines and their receptors that may be differentially modulated in these two T cell activation models and study the effects of SOS. In addition, stimulation of purified T cell subsets and surface/intracellular staining will help identify the cellular source of the cells responsible for producing these differentially produced cytokines/chemokines and their receptors. Additionally, the in vitro system does not capture the complete complexity of physiological T-cell activation, which occurs in three-dimensional lymphoid tissue environments with spatially organised antigen presentation, cytokine gradients and stromal cell interactions (Joseph *et al*., 2023, 2026). Finally, we will need to extend our study to human T cells to decipher the role of PKC signalling during human T cell activation.

In summary, TCR-independent activation with P+I and TCR-dependent activation with αCD3+αCD28 generate functionally and metabolically distinct T-cell responses. P+I showed superior proliferation, metabolic activity and activation-marker expression, and produced a distinct cytokine profile. The mechanistic basis for this disparity is insufficient PKC pathway engagement in the αCD3+αCD28 system, as demonstrated by direct PKC activity measurement (Figure 7) and confirmed by the selective rescue of proliferative, metabolic and phenotypic deficits by PMA, but not Ionomycin, supplementation (Figure 8, 9, 10, 11). These findings call for caution when comparing experimental results across the two activation platforms and suggest that augmenting PKC signalling in TCR-dependent activation is a rational and testable strategy for improving T-cell expansion in therapeutic contexts.

## Materials and Methods

### Reagents and Antibodies

Phorbol 12-myristate 13-acetate (PMA; Cat. No. P1585, Sigma-Aldrich), Ionomycin calcium salt from *Streptomyces conglobatus* (Cat. No. I0634, Sigma-Aldrich), RNase A (Cat. No. 19101, QIAGEN), propidium iodide (Cat. No. P4170, Sigma-Aldrich), DCFDA (Cat. No. D6883, Sigma-Aldrich), paraformaldehyde (Cat. No. P6148, Sigma-Aldrich), 2-NBDG (Cat. No. SML4177, Sigma-Aldrich), glucose-free RPMI-1640 (Cat. No. 11879020, Gibco, Thermo Fisher Scientific) and RPMI-1640 (Cat. No. AL060, HiMedia) were used in this study. The following fluorochrome-conjugated antibodies were used for flow cytometric analysis: PE-conjugated αCD4 (Cat. No. 100408, BioLegend), APC-conjugated αCD3 (Cat. No. 100236, BioLegend), Alexa Fluor 488-conjugated αCD8 (Cat. No. 100723, BioLegend), FITC-conjugated αCD69 (Cat. No. 11-0691-82, Invitrogen eBioscience), eFluor 450-conjugated αCD25 (Cat. No. 48-0253-82, Invitrogen eBioscience), PE-Cy5-conjugated αCD44 (Cat. No. 553135, BD Biosciences), and Spark Red 718-conjugated αPD-1 (Cat. No. 135260, BioLegend). For T cell isolation, goat anti-mouse IgG antibody used for panning was obtained from Jackson ImmunoResearch Laboratories, USA. For T cell activation, purified αCD3 (Cat. No. 16-0032-82, Invitrogen eBioscience) and αCD28 (Cat. No. 102101, BioLegend) antibodies were used.

### T cell isolation, activation and culture

All animal experiments were conducted in accordance with the guidelines of the Committee for the Purpose of Control and Supervision of Experiments on Animals (CPCSEA) and were approved by the Institutional Animal Ethics Committee (IAEC) of the Indian Institute of Science (IISc), Bengaluru, India (Approval No. CAF/Ethics/976/2023). T cells were isolated from mesenteric and inguinal lymph nodes of 6-8-week-old BALB/c male mice. Briefly, the lymph nodes were minced with forceps and passed through a 40 μm cell strainer. The single-cell suspension was prepared and incubated in a T25 flask pre-coated with 100 μg/mL goat anti-mouse IgG antibody for 30 min at 37°C to deplete B lymphocytes. This panning step was repeated to achieve improved purity of the T lymphocyte preparation. Purified T cells were stained for CD3 to confirm purity. The T cells were then seeded into U-bottom 96-well plates at a density of 5 × 10⁴ cells per well in RPMI 1640 medium supplemented with 5% FBS (Joseph *et al*., 2024). For TCR-independent activation, T cells were treated with P+IL (10 ng/mL PMA plus 0.2 μM Ionomycin) or P+IH (10 ng/mL PMA plus 0.8 μM Ionomycin). For TCR-dependent activation, 96-well U-bottom plates were coated with plate-bound αCD3 (0.5 μg/mL) with or without soluble αCD28 (1 μg/mL). For all experiments, cells were cultured at 37°C, 5% CO₂ and 95% humidity in 96-well U-bottom plates (Cat. No. 163320, ThermoFisher) (Joseph *et al*., 2024).

### Flow cytometry

Cell cycle analysis was performed using propidium iodide (PI) staining. Briefly, cells were fixed in 70% ethanol on ice for 15-30 min, washed with PBS, treated with 100 μg/mL RNase A for 30 min, then stained with 50 μg/mL PI in the dark before acquisition. For intracellular ROS estimation, cells were harvested at the 12 h and 24 h time points and stained with 20 μM DCFDA for 30 min prior to acquisition. To assess cell surface marker expression, antibodies were titrated against a range of concentrations prior to use: αCD3 (1:200), αCD4 (1:200), αCD8 (1:200), αCD69 (1:200), αCD25 (1:200), αCD44 (1:200) and αPD-1 (1:20). Antibodies were added to cells and incubated on ice for 30-45 min. Cells were then washed using PBS and fixed in 4% paraformaldehyde. The entire protocol was performed in the dark. After staining, cells were analyzed by flow cytometry. Live cell populations were gated using forward and side scatter plots, and MFI, percentage populations and histograms were plotted. All flow cytometry experiments were performed on CytoFLEX flow cytometer (Beckman Coulter, Brea, CA, USA) and data were analyzed using FlowJo v10.9 software (BD Biosciences, Ashland, OR, USA) (Joseph *et al*., 2024).

### Glucose uptake using 2-NBDG

T cells were first incubated in glucose-free RPMI for 1 h. Subsequently, 2-NBDG was added at a final concentration of 100 μM and then the cells were activated. The final measurement was taken following 2 h of activation based on timepoint titration (data not shown). Fluorescence intensity was measured at Ex: 465 nm, Em: 540 nm using a microplate reader (Infinite 200 PRO, Tecan Austria GmbH).

### Extracellular lactate estimation

Lactate estimation was performed using an L-Lactate Assay Kit (Cat. No. MAK329, Sigma). Cell culture supernatant was collected following T cell activation at 24 h and 36 h time points, and lactate concentration was determined according to the manufacturer’s protocol. The intensity of the coloured product is measured at 565 nm using a microplate reader which is proportional to the lactate concentration in the sample

### Estimation of cellular dehydrogenases activity

Cellular dehydrogenases activity was assessed using CCK-8 (Cat. No. 96992, Sigma). This assay is based on the reduction of the water-soluble tetrazolium salt (WST)-8 by cellular dehydrogenases in viable cells, producing a water soluble, orange coloured formazan dye whose intensity is measured at 450 nm using a microplate reader (Infinite 200 PRO, Tecan Austria GmbH). Cells were collected following T cell activation at 24 h and 36 h time points, and dehydrogenase activity was determined according to the manufacturer’s protocol.

### Cytokine and chemokine measurement

For cytokine and chemokine measurement, cell culture supernatants were collected at the 36 h time point and quantified by using eBioscience™ ELISA Ready-SET-Go™ kit, per manufacturer’s instructions.

### PKC activity measurements

PKC activity was quantified using an ELISA based PKC Kinase activity assay kit (Cat. No. ab139437; Abcam, Cambridge, UK) according to the manufacturer’s instructions. Equal numbers of T cells were stimulated with PMA + Ionomycin or αCD3 ± αCD28 antibodies for 30 min based on optimisation of time point (data not shown), followed by cell lysis using ice-cold lysis buffer containing 20 mM MOPS, 50 mM β-glycerophosphate, 50 mM sodium fluoride, 1 mM sodium orthovanadate, 5 mM EGTA, 2 mM EDTA, 1% NP-40, 1 mM DTT, 1 mM benzamidine, 1 mM PMSF, and 10 μg/mL each of leupeptin and aprotinin, as recommended by the manufacturer. Equal amounts of lysate were added to substrate coated wells in the presence of ATP to initiate the kinase reaction. Active PKC in the lysates phosphorylated the immobilised substrate peptide, which was detected using a phospho substrate specific primary antibody, followed by an HRP-conjugated secondary antibody. Colour was developed using tetramethylbenzidine substrate, and the reaction was stopped using stop solution. Absorbance was measured at 450 nm using a microplate reader. The absorbance reading was directly proportional to PKC activity in the samples.

### Apoptosis

Apoptosis was assessed using an Annexin-V FITC/PI staining according to the manufacturer’s protocol (cat. no. 640914; BioLegend, San Diego, CA, USA). Cells were resuspended in binding buffer and 100 μL was taken for staining. 5 μL Annexin-V FITC and 10 μL PI were added and incubated at room temperature in the dark for 15 min prior to acquisition. Data acquisition was done on CytoFLEX instrument, and data were analyzed using FlowJo software (Joseph *et al*., 2024).

### Statistical analysis

All data are presented as mean ± SEM from three independent experiments. Statistical significance was determined by one-way ANOVA with Tukey’s multiple comparisons test. P values are multiplicity adjusted; ns p > 0.05, *p < 0.05, **p < 0.01, ***p < 0.001, ****p < 0.0001. Graphs were generated using GraphPad Prism 10.

## Funding

This study was funded by a grant from the Indian Council of Medical Research (ICMR) EMDR/SG/11/2023-6777. In addition, the support by institute grants from IISc and infrastructural support from the FIST program of the Department of Science and Technology, India is greatly appreciated. NR appreciates the graduate student fellowship from the Department of Biotechnology DBT/2021-22/IISc/1555.

## Competing interests

The authors declare that they have no competing financial interests or personal relationships.

## Acknowledgements

The authors gratefully acknowledge the Divisional Flow Cytometry Facility and the Central Animal Facility, IISc. We thank all the members of the DpN lab for their support and encouragement. We are grateful to Dr. Payel Roy for insightful discussions and valuable inputs on this study. We also thank Micky Anand for critically reviewing the manuscript and for his constant support and encouragement throughout the course of this work. We thank the organisers of Immunocon-2026 and Molecular Immunology Forum-2026 for the opportunity to present this study and for the valuable feedback received during these meetings.

## Abbreviations

C: H ratio, Cycling-to-hypodiploidy ratio
I: Ionomycin
IH: Ionomycin high concentration
IL: Ionomycin low concentration
P: PMA
P+IL: PMA + Ionomycin (low)
P+IH: PMA + Ionomycin (high)
UA: Unactivated

## Notes

### Competing Interest Statement

The authors have declared no competing interest.

