## Supplemental Information for "TCR-dependent and TCR-independent *in-vitro* T cell activation generate distinct functional, metabolic, and cytokine programs: Protein kinase C signalling augments anti-CD3+anti-CD28 responses"

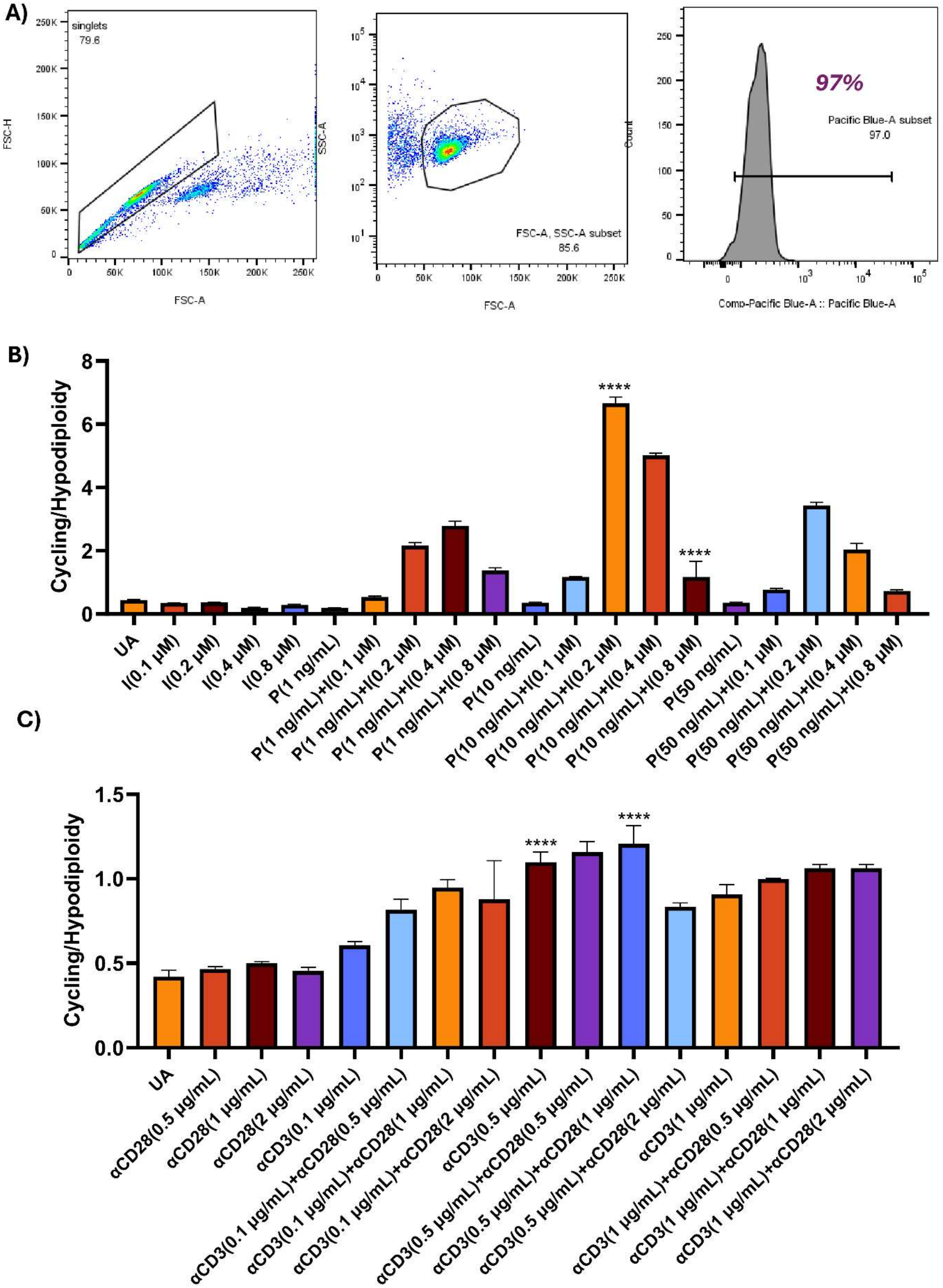
A)
singlets
79.6
FSC-H
FSC-A
250K
200K
150K
100K
50K
0
50K
100K
150K
200K
250K
SSC-A
FSC-A, SSC-A subset
85.6
Count
97%
Pacific Blue-A subset
97.0
Comp-Pacific Blue-A :: Pacific Blue-A
B)
Cycling/Hypodiploidy
8
6
4
2
0
****
****
UA
I(0.1 μM)
I(0.2 μM)
I(0.4 μM)
I(0.8 μM)
P(1 ng/mL)
P(1 ng/mL)+I(0.1 μM)
P(1 ng/mL)+I(0.2 μM)
P(1 ng/mL)+I(0.4 μM)
P(1 ng/mL)+I(0.8 μM)
P(10 ng/mL)
P(10 ng/mL)+I(0.1 μM)
P(10 ng/mL)+I(0.2 μM)
P(10 ng/mL)+I(0.4 μM)
P(10 ng/mL)+I(0.8 μM)
P(50 ng/mL)
P(50 ng/mL)+I(0.1 μM)
P(50 ng/mL)+I(0.2 μM)
P(50 ng/mL)+I(0.4 μM)
P(50 ng/mL)+I(0.8 μM)
C)
Cycling/Hypodiploidy
1.5
1.0
0.5
0.0
****
****
UA
αCD28(0.5 μg/mL)
αCD28(1 μg/mL)
αCD28(2 μg/mL)
αCD3(0.1 μg/mL)
αCD3(0.1 μg/mL)+αCD28(0.5 μg/mL)
αCD3(0.1 μg/mL)+αCD28(1 μg/mL)
αCD3(0.1 μg/mL)+αCD28(2 μg/mL)
αCD3(0.5 μg/mL)
αCD3(0.5 μg/mL)+αCD28(0.5 μg/mL)
αCD3(0.5 μg/mL)+αCD28(1 μg/mL)
αCD3(0.5 μg/mL)+αCD28(2 μg/mL)
αCD3(1 μg/mL)
αCD3(1 μg/mL)+αCD28(0.5 μg/mL)
αCD3(1 μg/mL)+αCD28(1 μg/mL)
αCD3(1 μg/mL)+αCD28(2 μg/mL)

**Supplementary Figure 1. Dose titration for TCR-independent (PMA+Ionomycin) and TCR-dependent (plate-bound αCD3 + αCD28) activation systems. (A)** Representative flow cytometry plots showing percentage of $CD3^+$ T cells after two rounds of panning, confirming T cell purity. **(B)** Cycling-to-hypodiploidy ratio for different combinations of PMA and Ionomycin. **(C)** Cycling-to-hypodiploidy ratio for different combinations of plate-bound αCD3 and soluble αCD28. Data are represented as mean ± SEM from three independent experiments Statistical significance was determined by one-way ANOVA with Tukey's multiple comparisons test. P values are multiplicity adjusted; ns $p > 0.05$, *$p < 0.05$, **$p < 0.01$, ***$p < 0.001$, ****$p < 0.0001$. **Abbreviations:** UA, unactivated; P, PMA; I, Ionomycin.

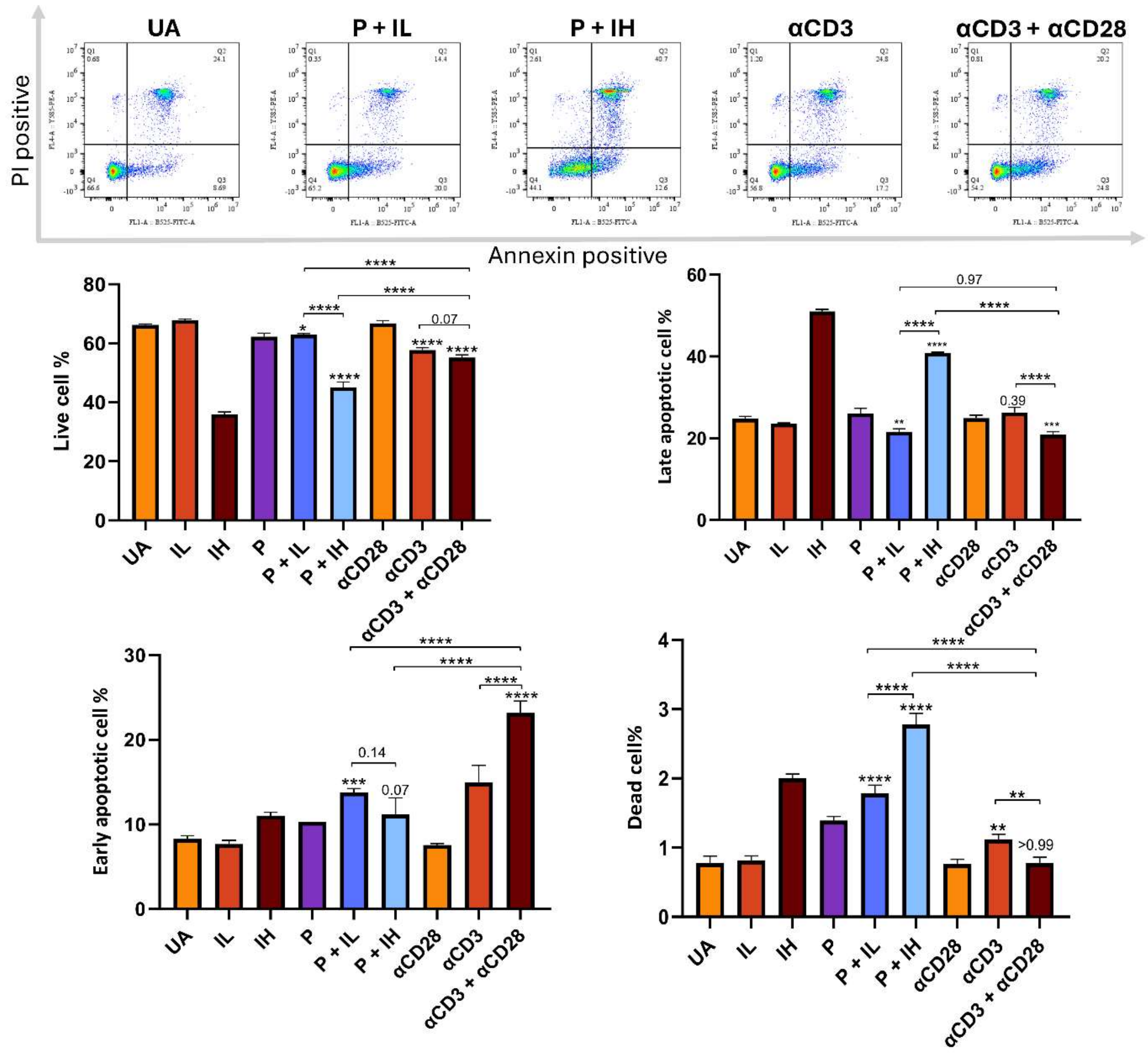


**Supplementary Figure 2. Assessment of apoptosis following T cell activation by Annexin V/PI staining.** T cells were activated under the indicated conditions and stained with Annexin V and PI at 36 h post-activation. Data are represented as mean ± SEM from three independent experiments Statistical significance was determined by one-way ANOVA with Tukey's multiple comparisons test. P values are multiplicity adjusted; ns $p > 0.05$, *$p < 0.05$, **$p < 0.01$, ***$p < 0.001$, ****$p < 0.0001$.

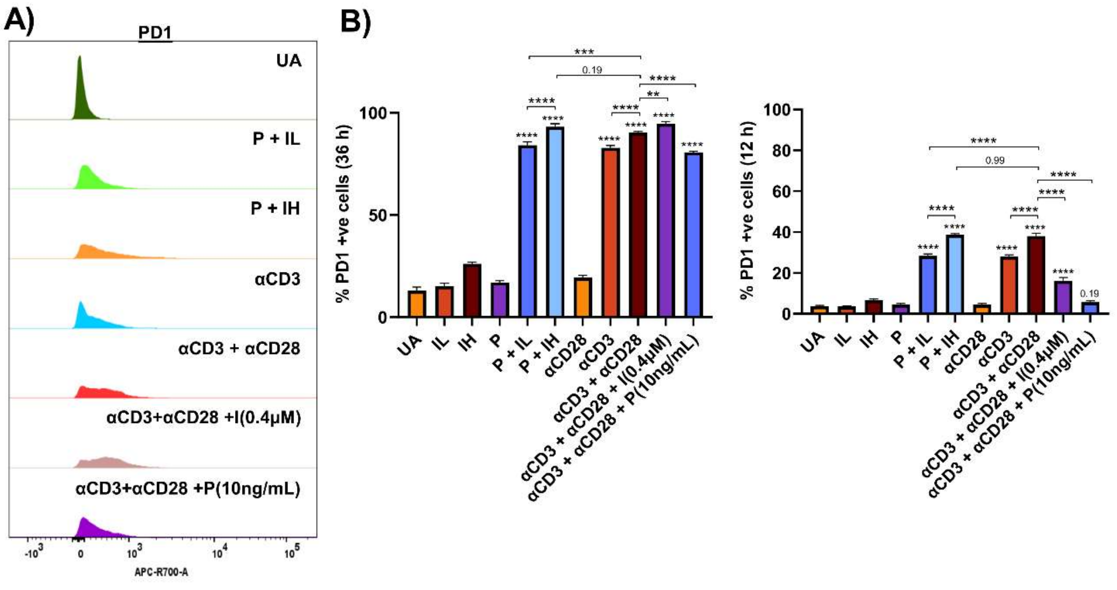


**Supplementary Figure 3. αCD3 + αCD28 stimulation leads to higher expression PD1 compared to P + IL and PMA supplementation lowers PD1 expression in αCD3 + αCD28 system. (A** Representative flow cytometric histograms showing surface expression of PD1 post-activation. **(B)** Fold change in MFI of PD1 post-activation. All data are normalised to unactivated control. Data are represented as mean ± SEM from three independent experiments Statistical significance was determined by one-way ANOVA with Tukey's multiple comparisons test. P values are multiplicity adjusted; ns $p > 0.05$, *$p < 0.05$, **$p < 0.01$, ***$p < 0.001$, ****$p < 0.0001$.
